# Real-time electrochemical protein monitoring using molecular pendulum sensors

**DOI:** 10.64898/2026.07.28.730971

**Authors:** Vuslat B. Juska, Zhiyuan Chen, Raza Qazi, Longshun Li, Saehyun Kim, Fatemeh Esmaeili, Will Buchsbaum, Audrey N. Nashner, Jane M. Donnelly, Luis Fernando Ayala-Cardona, Ryan A. Neff, Andrew J. H. Sedlack, Maria D. Cabezas, Jagotamoy Das, Shana O. Kelley, Hossein Zargartalebi

## Abstract

Continuous monitoring of proteins in complex biological fluids is essential for advancing personalized medicine, yet existing biosensors are often limited by instability, single-use designs, and insufficient sensitivity. Here, we describe a detailed protocol for the fabrication and operation of molecular pendulum (MP) electrochemical sensors integrated with an active-reset mechanism to enable real-time, reversible, and ultrasensitive protein detection. The protocol is described in two parts. First, we describe the microfabrication of gold microelectrodes and their nanostructured modification, followed by assembly of DNA-based pendulum probes with redox reporters and affinity receptors that translate binding into kinetic electron-transfer shifts. Second, we detail the sensing and active-reset approach, which detects the target analyte and applies tunable oscillatory potentials to accelerate its dissociation, regenerate sensor surfaces, and extend operational lifetime. The protocol includes detailed guidance on device fabrication, surface functionalization, sensing and reset cycles, and data analysis. When implemented, MP sensors with active-reset achieve pg/ml sensitivity, rapid equilibration, and robust performance across biofluids and in situ models, enabling continuous protein monitoring over extended periods. This combined technology represents a biosensing platform with significant potential, opening new avenues for wearable and implantable molecular monitoring, early disease detection, and personalized therapeutic guidance.

## Introduction

### Background

Continuous monitoring of protein and small-molecule biomarkers is essential for understanding dynamic biological processes such as inflammation, immune signaling, and tissue injury, which often evolve on timescales that are inaccessible to conventional analytical methods^1–5^. Moreover, key inflammatory protein biomarkers, including interleukin-6 (IL-6) and tumor necrosis factor-α (TNF-α), fluctuate rapidly and exhibit short effective half-lives in vivo, rendering single-time-point measurements ill-equipped to inform clinical actions^6^. Most established protein detection techniques rely on discrete sampling, reagent addition, and multistep workflows, which fundamentally limit temporal resolution and preclude long-term measurements in living systems^7^.

Reagentless and continuous sensing strategies address these limitations by embedding molecular recognition directly into the sensor architecture, enabling repeated measurements without consumables or sample processing^8–11^. Among these approaches, electrochemical sensors are particularly attractive due to their scalability, compatibility with miniaturized electronics, and robust operation in complex and optically opaque biological environments, making them well suited for wearable and in vivo applications.

### Development of molecular pendulum sensing

MP sensing is a DNA-based, reagentless electrochemical approach that converts protein binding events into changes in electron-transfer kinetics through electrically driven molecular motion^8,12,13^ (Fig. 1a). In this architecture, the molecular pendulum scaffold is formed by hybridization of two negatively charged DNA strands, Probe 1 (P1) and Probe 2 (P2). P1 is tethered to the gold electrode surface through a terminal thiol anchor and carries a ferrocene redox reporter at its distal end. P2 hybridizes to the complementary region of P1 and carries the molecular recognition element, such as an antibody, aptamer or nanobody. Together, the P1–P2 duplex forms the surface-confined pendulum structure that links target binding to changes in electrochemical electron-transfer kinetics ^8,12,14^.

**Fig. 1.**
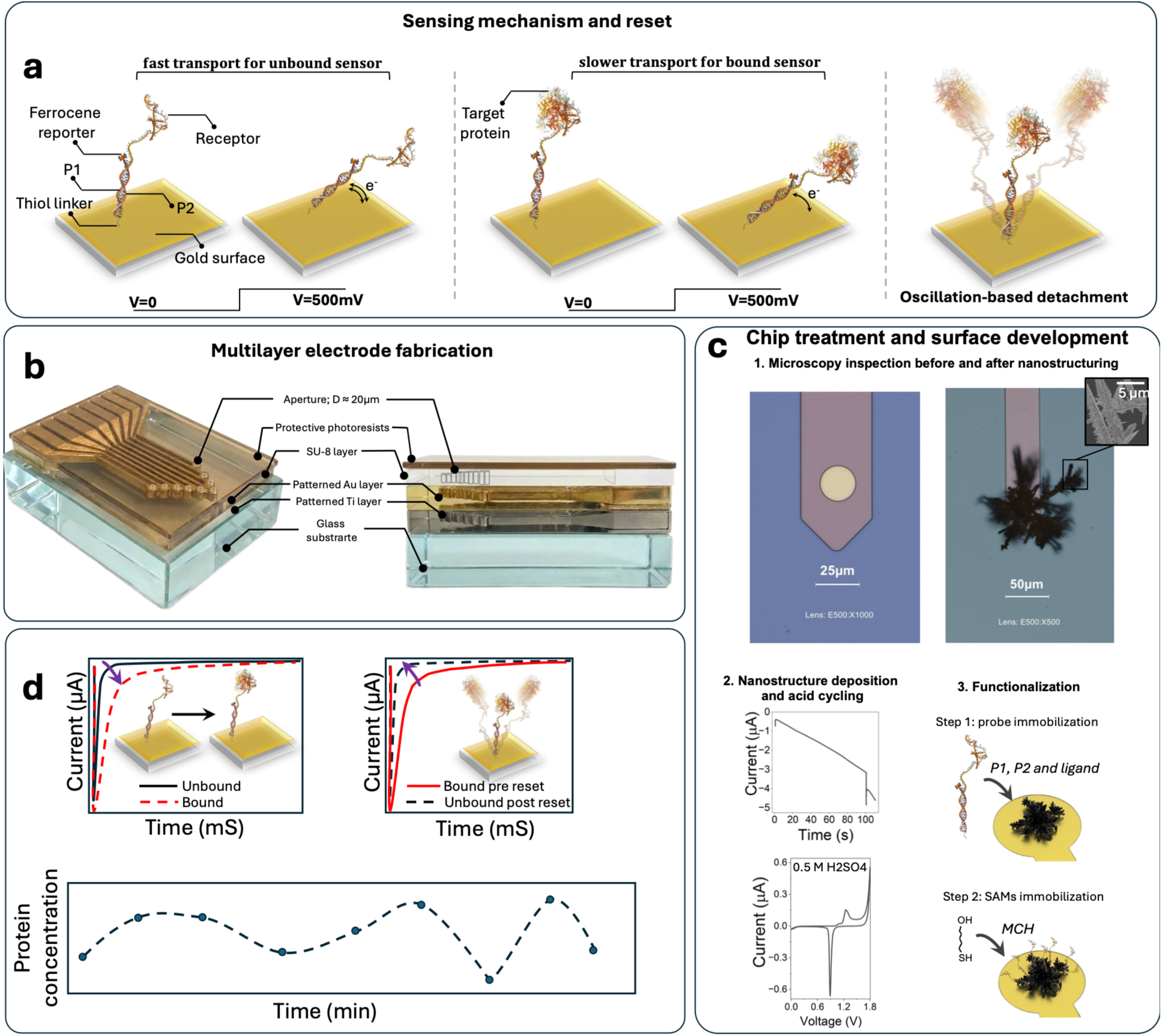
Overview of the real time protein monitoring protocol. **a,** molecular pendulum (MP) architecture, sensing mechanism, and active-reset. **b,** Multilayer electrode used for MP sensing (Steps 1-21). Schematics were generated with AI assistance and are not drawn to scale; all scientific content was reviewed by the authors for accuracy. **c,** Chip preparation and functionalization procedure (Steps 43-45). **d,** MP sensing and active-reset lead to continuous protein monitoring (Steps 46-55).

Upon application of a positive electrode potential, the electric field drives the negatively charged DNA construct toward the electrode surface. When the applied potential exceeds the oxidation potential of ferrocene (approximately 380 mV vs Ag/AgCl), electron transfer occurs between the redox reporter and the electrode, generating a measurable faradaic current. Binding of the target protein to the recognition element increases the hydrodynamic drag experienced by the DNA construct, leading to reproducible changes in the rate and magnitude of electron transfer during chronoamperometric interrogation (Fig. 1a).

### Comparison with conventional electrochemical affinity sensors

The electrically actuated nature of MP sensing fundamentally distinguishes it from conventional structure-switching electrochemical affinity sensors, in which target binding induces a conformational rearrangement of the probe that modulates the average proximity of a redox reporter to the electrode surface^10^. In these structure-switching architectures, signal transduction requires that the receptor, most commonly an aptamer, undergo a sufficiently large and reproducible binding-induced conformational change; in the absence of such a transition, the sensor produces little or no useful signal change. Consequently, the applicability of this sensing modality is largely confined to receptors that intrinsically support target-coupled structural switching, thereby restricting the range of compatible aptamers and limiting broader receptor generality. In such systems, signal generation is governed by equilibrium structural changes, limiting temporal control over sensor interrogation^11^. By contrast, MP sensing decouples signal transduction from passive probe fluctuations by using an applied electric field to actively drive probe motion.

Because the sensing signal in MP sensors is encoded in electron-transfer kinetics rather than solely in equilibrium binding, this approach is well suited for high-frequency, time-resolved measurements. This capability is particularly advantageous for monitoring protein dynamics in vivo, where concentrations can fluctuate rapidly and transient signaling events are common^2^. In addition, the modular design of the P2 domain enables straightforward substitution of different receptor classes without altering the underlying transduction mechanism, allowing adaptation of the platform to a broad range of protein targets^13^.

### Sensitivity of MP sensors

MP sensors reproducibly detect analytes at concentrations often several orders of magnitude below the solution-phase K_D_ of the receptors employed. Our work with this approach includes robust control strategies and careful drift monitoring to validate that a response is truly analyte-specific^15^. The observation of detection limits significantly below those predicted by solution-phase K_D_ values is shared across broad classes of affinity-based sensors, and it is therefore clear that a simple, solution-phase thermodynamic model is not sufficient to describe the behavior of these systems^16–27^. Equilibrium dissociation constants quantify receptor-target affinity under specific conditions that differ from sensing trials, and carry no information about surface confinement, probe density, interfacial electrostatics, local rebinding, or the kinetic selectivity of time-resolved electrochemical readout^27^. Each of these factors operates independently of the solution-phase thermodynamics and contributes to sensitivity in ways that a K_D_-based model cannot anticipate. The appropriate framework for understanding electrochemical affinity sensor performance is therefore not equilibrium thermodynamics alone, but a kinetic and interfacial description that treats the sensing surface as a physicochemically distinct environment from the bulk solution in which receptor affinities are conventionally measured.

The MP architecture provides several distinct and reinforcing physical mechanisms that collectively account for this sensitivity without invoking drift or artifact. First, the solution-phase K_D_ measured by biolayer interferometry or surface plasmon resonance is not necessarily the operative affinity constant at the sensing interface. Nanostructuring of the gold electrode surface – a key feature of the overall sensing platform – creates a high-curvature, high-density probe environment in which electrostatic pre-concentration, altered probe orientation, and cooperative neighbor effects can substantially lower the effective surface-phase dissociation constant relative to its solution-phase value^28–30^. The K_D_ values provided in this protocol should therefore be understood as conservative upper bounds on the affinity experienced at the nanostructured electrode surface.

Second, surface confinement of both the probe and the target may suppress three-dimensional diffusion away from the sensing layer. A protein that dissociates from one probe encounters neighboring probes before escaping to bulk solution, increasing effective dwell time and instantaneous fractional occupancy beyond what solution-phase kinetics would predict. This local rebinding effect means that even receptors with fast k_off_ can maintain a disproportionately high instantaneous occupied fraction at the surface.

Third, the per-event signal change in MP sensing appears to be intrinsically large. Signal transduction depends on perturbation of the hydrodynamic drag experienced by the DNA construct during electrically driven motion, and this drag is not determined solely by the dry molecular mass of the bound protein. In solution, the effective hydrodynamic size of a protein reflects its shape, surface chemistry, conformational state and coupling to surrounding solvent, including dynamic hydration and entrained water associated with the protein surface^31^. As a result, the hydrodynamic perturbation introduced by target binding can exceed that expected from the anhydrous protein volume alone. This solvent-coupled increase in effective hydrodynamic load may amplify the drag perturbation per binding event relative to the unloaded pendulum arm, contributing to high transduction gain even at low fractional occupancy.

Fourth, because chronoamperometry (CA) is a time-resolved kinetic measurement rather than an equilibrium readout, the sensor may be sensitive to transient molecular encounters that would be invisible to ensemble affinity methods. At sub-K_D_ concentrations, the time-averaged occupancy is low, but the instantaneous encounter rate across the probe ensemble can be sufficient to ensure that a meaningful fraction of pendulum molecules is transiently loaded at any moment of interrogation. Encounter complexes with dwell times comparable to the CA measurement window contribute real drag perturbations to the recorded signal, regardless of whether they constitute stable binding by equilibrium criteria. This sensitivity to transient interactions is a fundamental distinction from structure-switching sensors, which require a stable receptor conformational change and are therefore blind to short-lived encounters.

Fifth, and perhaps most importantly, the CA readout provides inherent background suppression through a kinetic gating mechanism. Free, unbound pendulum probes execute rapid field-driven motion and complete their transit to the electrode within a characteristic time window that defines the analytical measurement point. Only pendulum probes that exhibit target binding produce the delayed transit signature that constitutes the analytical signal. This self-filtering property means that the MP readout is intrinsically selective for the bound state without requiring physical separation, washing, or signal correction, and it provides a degree of background suppression against both non-specific adsorption and surface fouling that is not available to equilibrium-based electrochemical sensors.

Taken together, these mechanisms, surface-phase affinity enhancement, local rebinding at the confined interface, large per-event hydrodynamic perturbation including hydration shell contributions, sensitivity to transient encounters through time-resolved interrogation, and kinetic gating against non-specific background, provide a coherent physical framework for sub-equilibrium sensitivity that is grounded in well-established principles of surface chemistry, hydrodynamics, and electrochemical kinetics. Developing a quantitative framework will require a concerted effort bringing together the needed expertise across these fields.

### Limitations

A central challenge in extending DNA-based electrochemical sensors to continuous operation is maintaining signal stability over prolonged measurement periods in complex biological environments^15,32,33^. Nonspecific adsorption, surface fouling, and gradual reorganization of the sensing interface can lead to baseline drift, attenuated signal amplitude, and reduced measurement fidelity^34–36^. Although these effects can be partially mitigated through optimization of self-assembled monolayers or addressed through post hoc signal correction, such approaches do not fully resolve instability during long-term operation.

In addition to biofouling and affinity-limited reversibility, MP sensing imposes specific electrochemical design constraints related to signal transduction. Because sensing relies on resolving binding-induced changes in faradaic current kinetics, the non-faradaic (capacitive) background current must be sufficiently low to enable reliable discrimination of the faradaic response. This requirement places strict constraints on the geometric surface area of the working electrode. In practice, effective operation of MP sensors requires microelectrodes with micron-scale apertures, where capacitive charging currents are minimized while maintaining adequate faradaic signal amplitude. Macroelectrodes that have capacitive decay characteristics extending into the window of the faradaic pendulum response are incompatible with this sensing approach^37^. Given the kinetic nature of the readout for MP sensors, minimization of capacitive current is critical and therefore, the optimal output can be achieved when the electroactive surface area is maintained at microscale dimensions (see Box 1). At present, most commercially available electrode arrays do not provide the requisite surface area control and reproducibility needed for MP operation. Moreover, a high-quality gold surface lacking any competing electrochemical signals is required. For example, in our experience, printed circuit boards or screen printed arrays with patterned gold electrodes do not provide the gold surface quality or precisely controlled electroactive area required for MP sensing. As a result, implementation of the platform necessitates custom microfabrication, typically involving cleanroom-based photolithographic patterning of gold microelectrodes or alternative strategies to precisely confine the effective working electrode area to the micrometer scale. Consequently, the fabrication workflow described in this protocol is a critical determinant of sensor performance, stability, and reproducibility and represents a practical barrier to adoption in settings without access to microfabrication facilities.

A further limitation of conventional affinity-based sensors arises from the requirement to detect low-abundance protein targets in vivo using high-affinity receptors, which frequently exhibit slow dissociation kinetics^38,39^. Prolonged target residence times hinder sensor regeneration and preclude truly continuous measurements. An active-reset strategy has therefore emerged as a universal solution to affinity-limited reversibility by enabling periodic electrochemical reconditioning of the sensing interface. For DNA-based electrochemical sensors, active reset typically involves the application of controlled oscillatory voltage waveforms that induce probe motion, accelerate target dissociation, and restore sensor responsiveness^2^ (Fig. 1a). These electrical perturbations simultaneously disrupt nonspecific interactions and reestablish reproducible DNA-electrode coupling without device removal or reagent replenishment. When integrated with MP sensing, active-reset operation enables sustained real-time protein monitoring in vivo^2^. Alternative approaches are also emerging that utilize simpler protocols focused on non-oscillating fields^40–42^.

### Applications and scope of the protocol

The MP sensing platform described in this protocol is designed for applications that require continuous, high-temporal-resolution monitoring of protein biomarkers in complex biological environments. In particular, the method is well suited for tracking dynamic inflammatory and immune signaling processes in vivo, where protein concentrations can fluctuate rapidly and transient events may be missed by intermittent sampling approaches. Representative applications include real-time monitoring of cytokines and inflammatory mediators during acute immune responses, tissue injury, infection, and therapeutic intervention^43^.

Beyond inflammatory biomarkers, the modular architecture of the MP enables adaptation of the platform to a wide range of protein targets by straightforward substitution of the recognition element, including antibodies, aptamers, and nanobodies. This flexibility supports applications in longitudinal pharmacodynamic monitoring, assessment of disease progression, and evaluation of treatment response, particularly in settings where repeated blood draws or ex vivo analysis are impractical or infeasible.

This protocol provides a comprehensive workflow for fabricating and operating MP sensors with active-reset capability, enabling sustained real-time protein monitoring over extended periods. While the methods described here are demonstrated using microfabricated gold electrodes within a controlled laboratory environment, the underlying principles are compatible with integration into wearable and implantable devices for continuous molecular monitoring in preclinical and translational settings. This protocol is recommended for those with significant experience in electrochemical sensor development and testing, microfabrication, as well as handling of sensitive protein analytes that are prone to degradation.

### Experimental design

The experimental design for real-time protein monitoring using MP sensors comprises four tightly linked stages: device fabrication, chip preparation and surface functionalization, analyte sensing and calibration, and active-reset-enabled continuous monitoring (Fig. 1). Each stage must be carefully controlled, because the overall performance of the platform depends not only on the quality of the sensing chemistry but also on electrode reproducibility, surface passivation, biomolecular assembly, electrochemical operating conditions, and the temporal fidelity of the instrumentation used to interrogate the sensor. Small deviations introduced at any stage can propagate through the workflow and degrade sensor sensitivity, reproducibility, baseline stability, transient fidelity, or resetting efficiency.

### Device fabrication

The first stage involves fabrication of the microelectrode chip by cleanroom-based photolithography and soft lithography (Figs. 1b and 2). This stage establishes the physical platform on which MP sensing is performed and is therefore critical for achieving reproducible electrochemical performance across devices. Patterned Ti/Au electrodes on glass, together with an SU-8-defined aperture, provide a well-confined sensing area and a multilayer architecture suitable for electrochemical interrogation. Particular attention should be paid to substrate cleanliness, photoresist processing, metal etching or lift-off quality, and SU-8 pattern fidelity, as these directly affect electrode geometry, exposed surface area, passivation integrity, and eventual sensor-to-sensor variability. Microscopic inspection of the patterned electrodes and apertures should be used routinely to verify structural integrity before downstream functionalization (Fig. 2b,c). If needed, topographical and morphological characterization methods (e.g., profilometry, atomic force microscopy (AFM) and scanning electron microscopy (SEM)) together with electrochemistry can be incorporated as quality-control steps to confirm etch-depth, aperture dimensions, metal continuity, and bare electrode behavior before biomolecular modification.

**Fig. 2.**
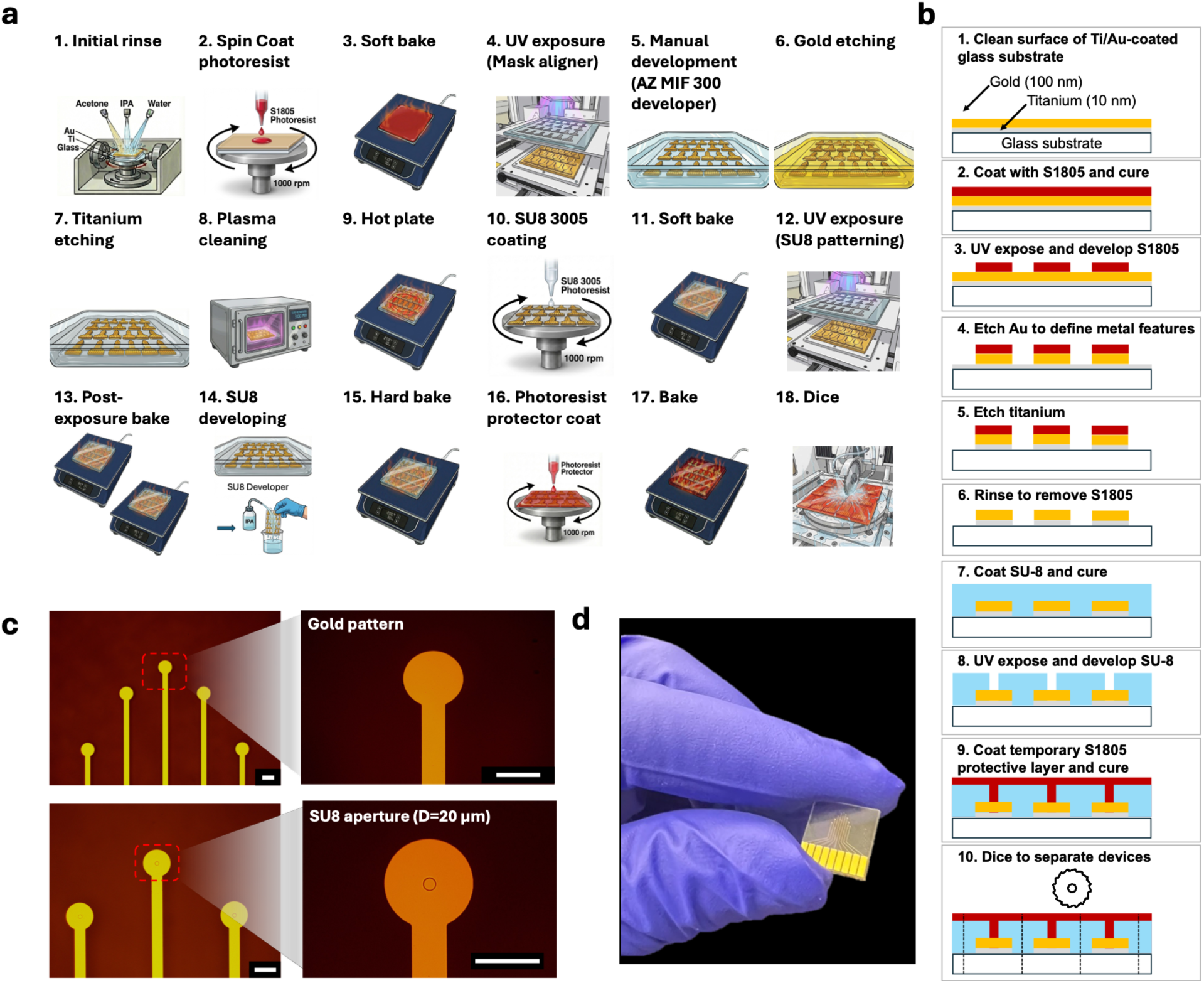
Cleanroom-based chip fabrication by soft lithography. **a,** Detailed workflow for patterning glass-based gold electrodes (steps 1–31). Schematics were generated with AI and author verified. **b,** Step-by-step side-view schematic illustrating the electrode microfabrication process**. c,** Microscopy image of the patterned gold electrode and SU-8 aperture with a 20 µm diameter; scale bars: 100µm. **d,** Image of a fabricated glass-based chip containing nine patterned gold working electrodes.

### Chip preparation and surface functionalization

The second stage consists of chip preparation and surface functionalization to assemble the MP architecture on the electrode surface (Fig. 1c). This step defines the sensing interface and is the core determinant of molecular recognition, signal transduction, and operational stability. The immobilized MP construct must be assembled in a manner that yields sufficient surface density for measurable signal generation while preserving enough conformational freedom for field-driven motion and target-responsive behavior. Surface blocking and passivation conditions should be optimized to suppress nonspecific adsorption and background drift without restricting probe mobility. The design of the sensing layer, including linker structure, redox reporter positioning, probe orientation, and spacing at the electrode interface, should be considered carefully because these factors influence electron-transfer efficiency, hydrodynamic drag, target accessibility, and the extent to which the bound and unbound states can be resolved electrochemically. Because surface chemistry is often a major source of batch-to-batch variability, functionalization should be performed under highly controlled conditions, and sensors should be equilibrated and screened before use.

### Analyte sensing and calibration

The third stage is analyte sensing and calibration, in which the electrochemical response of the MP sensor is measured as a function of target concentration (Figs. 1d and 3). In this protocol, CA interrogation is used to monitor the state-dependent motion of the pendulum under an applied electric field, thereby converting biomolecular binding into a measurable time-resolved current response.

**Fig. 3.**
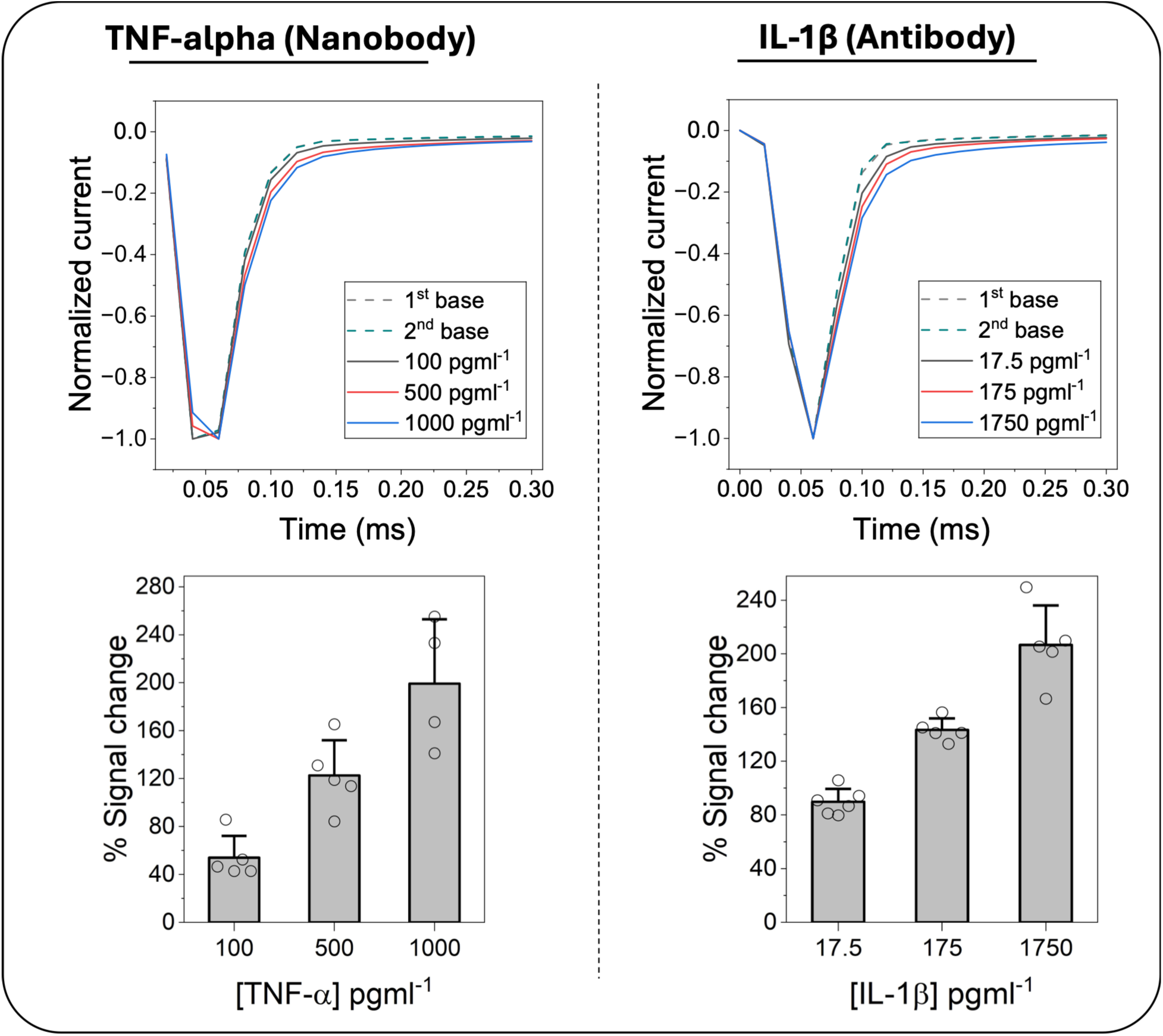
MP sensing using different biorecognition elements. Calibration curves for nanobody receptor detecting Tumor Necrosis Factor-alpha (TNF-α) and antibody receptor detecting interleukin-1β (IL-1β), together with the corresponding raw CA signals. Bar graphs show mean values, with error bars representing the standard deviation.

### Instrumentation considerations for high-speed MP sensing

Because MP sensing relies on rapid, field-driven motion of a surface-confined redox-active construct, the fidelity of the measured CA response depends not only on the sensor architecture and surface chemistry but also on the performance of the instrumentation used to interrogate it. In this regime, the applied potential waveform, control-loop bandwidth, transimpedance stage, sampling interval, and overall settling behavior of the potentiostat can directly shape the recorded transient, potentially obscuring or distorting the underlying molecular response. As a result, instrumentation is not merely an implementation detail for MP sensing, but a critical determinant of whether fast binding-dependent motion is captured faithfully or significantly distorted due to the convolution with the instrument’s response function. For this reason, high-speed electrochemical measurements used throughout this protocol should be performed with careful attention to potentiostat bandwidth and stability, measurement accuracy and fidelity, digitization sampling rate and quantization noise, wiring and shielding, and validation using model electrical loads when appropriate.

In fast CA measurements, the recorded signal represents the combined response of the electrochemical cell and the measurement hardware rather than the intrinsic sensor response alone. If the potentiostat bandwidth, rise time, or settling characteristics are not sufficiently faster than the transient of interest, sharp signal features may be attenuated, broadened, aliased, or replaced by instrument-limited behavior. This is particularly important for MP sensing, where analytical information is encoded in rapid current decays arising from target-dependent changes in pendulum motion. Accordingly, potentiostat selection should be guided not only by nominal sensitivity but also by control-loop bandwidth, applied waveform fidelity, transimpedance gain-bandwidth trade-offs, and output data rate needed for the experiment (see Box 2).

The digitization strategy should likewise be considered part of the experimental design. A digitizer that samples too slowly will miss or misrepresent the early portion of the transient, whereas aggressive post-acquisition smoothing can remove genuine kinetic information together with noise. Faster analog-to-digital conversion architectures and principled oversampling approaches are generally better suited than heavy post hoc filtering for preserving transient fidelity. However, prioritizing sampling speed must not come at the expense of vertical resolution. If the analog-to-digital converter (ADC) samples fast but lacks sufficient bit depth, quantization noise can easily overshadow the minute but critical signal changes in the measurement. When possible, the performance of the chosen instrument should be verified under the same operating conditions used for sensing rather than inferred from manufacturer specifications alone. Such tests provide a practical way to distinguish true sensing behavior from artifacts introduced by bandwidth limitations, phase-margin instability, or inadequate sampling.

The integrity of fast MP measurements also depends on proper control of electrochemical cell geometry and signal routing. In a three-electrode configuration, the working electrode potential is regulated relative to the reference electrode through a negative-feedback control loop, and any instability, excessive uncompensated electrolyte resistance, or poor reference-electrode placement can shift the true interfacial potential away from the intended value. Even when the nominal timescale is slower than that used in ultrafast electrochemistry, ohmic drop and control-loop limitations can still distort transient measurements if the electrolyte resistance is high or the reference electrode is positioned too far from the sensor, necessitating explicit uncompensated ohmic drop compensation to correct the dynamically shifting potential. The current generated at the electrode is converted to voltage by the transimpedance stage, and at the picoampere-to-microampere levels relevant to surface-confined protein sensing, stray capacitance, unshielded connections, and environmental electromagnetic interference can substantially elevate noise or compromise stability. To preserve signal integrity, careful hardware design techniques are paramount: electrical paths must be kept short, grounding and shielding must be rigorously applied, and all wiring must be rigidly secured to prevent movement-induced noise, supplemented by Faraday-cage measurements where appropriate.

Calibration experiments with representative inflammatory proteins, including TNF-α and interleukin-1 beta (IL-1β), establish the dynamic range, analytical sensitivity, and reproducibility of the sensing platform (Fig. 3). Experimental design at this stage should account for the large differences in molecular size, charge distribution, binding kinetics, and transport behavior among protein analytes, as these can strongly influence MP motion and signal magnitude. Appropriate buffer composition, temperature control, equilibration time, and replicate measurements are required to ensure that observed signal changes arise from target recognition rather than from nonspecific interfacial changes or instrumental drift. Extreme care must be taken with protein analytes tested at low concentrations to avoid degradation or adsorption to vessel surfaces.

### Active-reset-enabled continuous protein monitoring

The final stage is active-reset-enabled continuous protein monitoring (Figs. 1d and 4), which distinguishes this platform from conventional affinity-based biosensors limited by slow dissociation kinetics. In this stage, an electrical oscillation waveform is applied to actively accelerate target dissociation and regenerate the sensing interface, thereby enabling repeated cycles of measurement and reset on a single device. Successful implementation of this step requires careful tuning of reset parameters, especially voltage amplitude, frequency, waveform timing, and the balance between detection and regeneration periods. These parameters must be chosen such that resetting is efficient enough to prevent cumulative occupancy of the sensing layer, while avoiding damage to the surface chemistry, perturbation of the monolayer, or excessive baseline distortion. Continuous-monitoring experiments should therefore be designed to evaluate not only signal recovery after reset, but also signal stability, cycle-to-cycle reproducibility, transient fidelity, and resistance to progressive degradation over repeated operation. Monitoring IL-1β across changing concentrations provides a representative example of how the MP sensing strategy can be combined with active-reset to track dynamic protein changes over time while maintaining sensor responsiveness (Fig. 4).

**Fig. 4.**
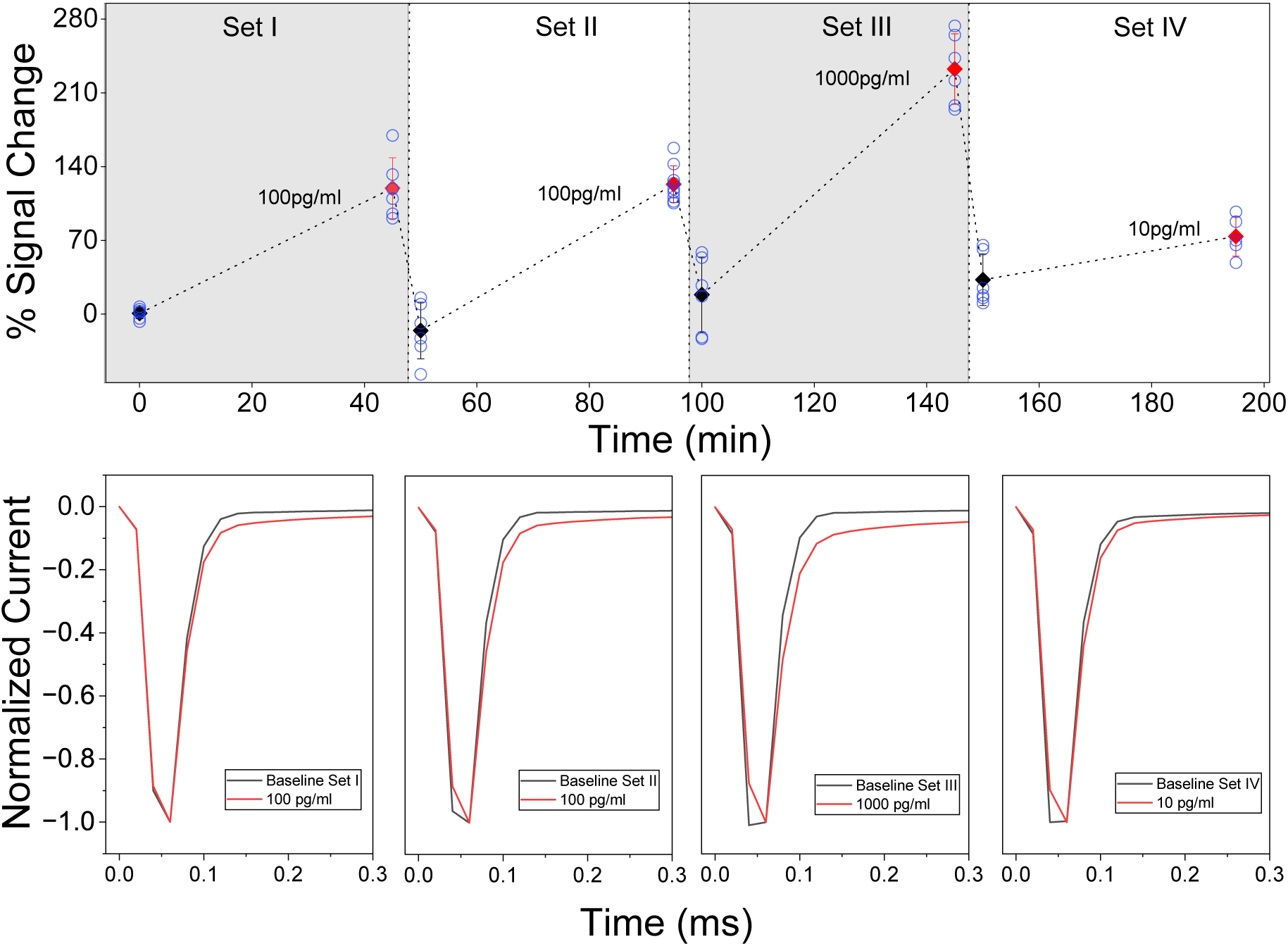
Active-resetting of the sensor for continuous protein monitoring. Continuous monitoring of IL-1β at different concentrations, together with raw CA traces showing detection and reset signals; error bars indicate the standard deviation.

### System-level considerations for reproducibility and optimization

Because the workflow integrates microfabrication, surface chemistry, electrochemical measurement, instrumentation, and kinetic control, reproducibility depends on treating the protocol as a unified system rather than a series of independent steps. Device geometry, surface assembly, electrochemical parameters, and hardware response are strongly coupled: for example, changes in aperture size or probe density can alter current magnitude and interfacial capacitance, while changes in waveform settings or instrument response can affect both target dissociation and the apparent transient shape. For this reason, optimization should proceed iteratively, with fabrication quality, surface functionalization, instrumentation performance, and sensing behavior evaluated together. When each stage is executed properly, the protocol yields a robust platform for real-time monitoring of protein dynamics and provides a general framework for extending electrochemical affinity sensing toward continuous in situ biomarker measurement.

#### BOX 1 Electrode surface area considerations

**Fig. 1.**
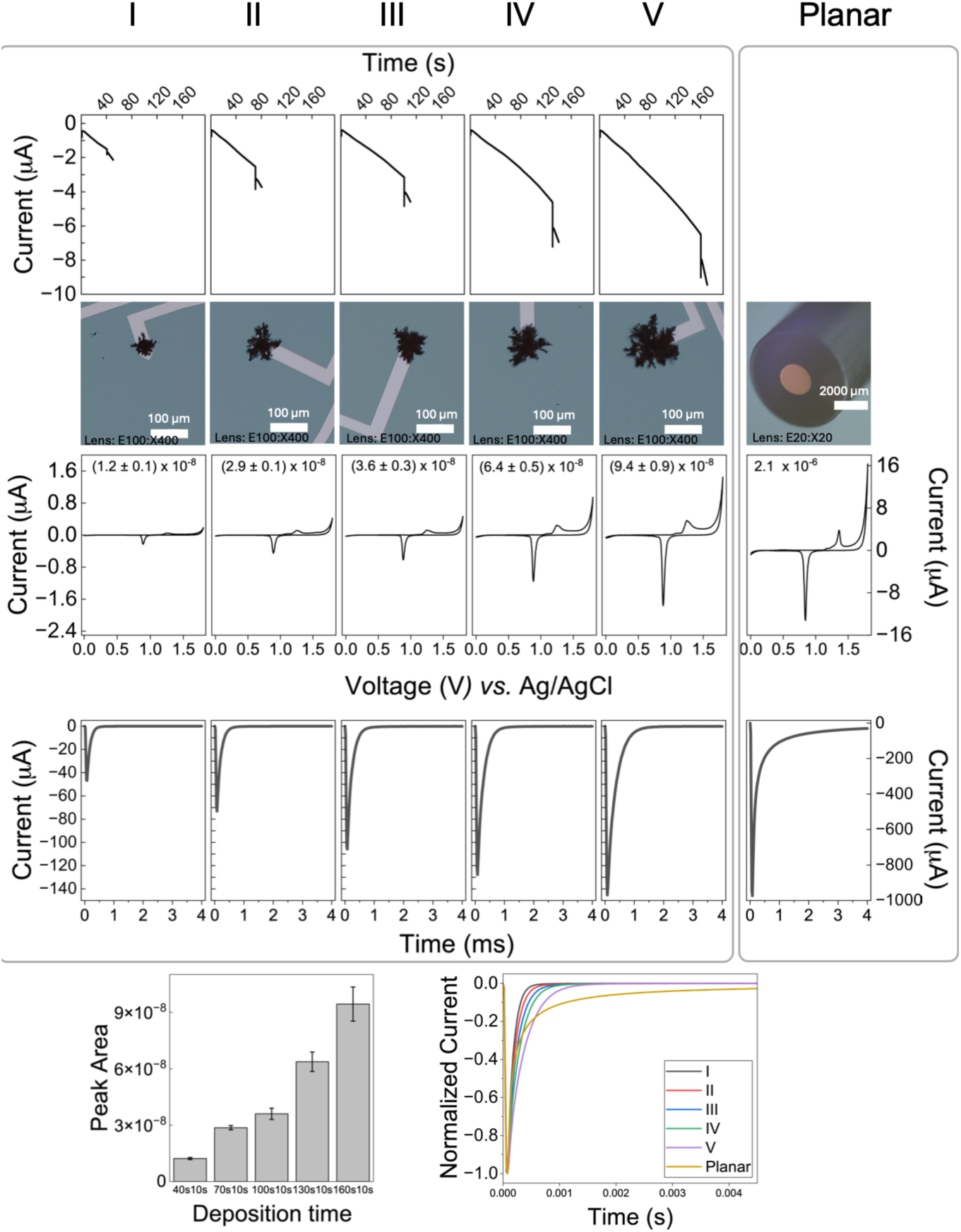
Comparison of capacitive current decays for microelectrodes and a planar macroelectrode. The small microelectrodes exhibit significantly faster current decays than the macroelectrode. For pendulum-based sensing, minimizing the capacitive current is desired to enable deconvolution from the faradaic current. The CVs were recorded in 0.5 M H₂SO₄ at a scan rate of 0.1 V s⁻¹, and the integrated gold oxide reduction peak area is reported in µA·V. Bar graphs show mean values, with error bars representing the standard deviation.

#### BOX 2 Effect of potentiostat acquisition capabilities on fast CA signal capture

To illustrate how potentiostat configuration affects fast CA measurements, we recorded a pair of fast current transients, current 1 and current 2, from two RC dummy circuits chosen to produce a small but resolvable difference in their decay (< 500nA), using a BASi Epsilon Eclipse potentiostat under several combinations of sampling interval and analog bandwidth (Fig. 1). An LTSpice simulation of the expected transient is included as an idealized reference for the current profile in the absence of instrumental bandwidth and digitization limits and provides an approximate source of truth for the peak signal amplitude. Across all settings, the instrument reproduced the overall shape of the transients and resolved the difference between current 1 and current 2, but the recovered signal amplitude and the fidelity of the earliest portion of the transient varied with the acquisition configuration, with some settings recovering only a fraction of the signal of interest. Because a fast-CA transient carries information in both its frequency content and its short time constants, an instrument must provide both adequate analog bandwidth and a sufficiently short sampling interval to capture it faithfully, and settings that are limiting in either respect will under-report the fast signal. The specific settings that best preserved the transient here reflect the time constants of this particular test signal and are not intended as general recommendations.

These results show that potentiostat parameters, including analog bandwidth, current-range response, filtering, digitization rate, and time resolution, can influence the measured shape and amplitude of fast CA transients and, therefore, the measurement of molecular pendulum responses. With a 20 µs sampling interval and 10kHz bandwidth, the BASi system can capture the difference between the two current transients in these measurements, although the earliest current features may be attenuated or broadened relative to the simulated response. Instruments or acquisition settings with lower temporal resolution may distort the transient more substantially, potentially reducing sensitivity to the fast signal components relevant to molecular pendulum measurements.

**Fig. 1.**
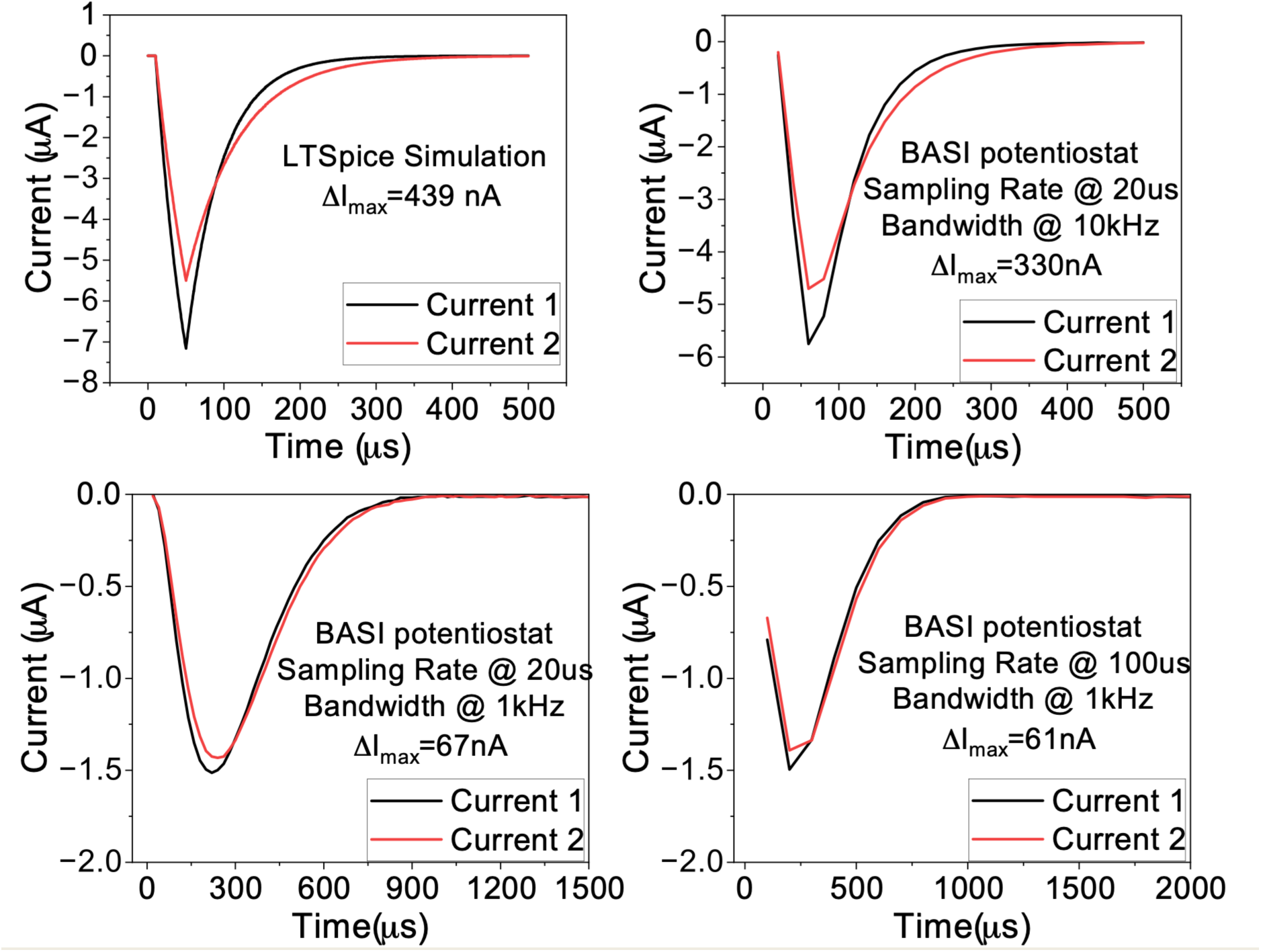
Effect of potentiostat bandwidth and sampling interval on fast chronoamperometric transients. Two RC dummy circuits, producing currents 1 and 2 mimicking an unbound and bound sensor respectively, were measured with a BASi Epsilon Eclipse potentiostat at different combinations of analog bandwidth and sampling interval, and compared with an LTSpice simulation. At a 20 µs sampling interval and 10 kHz bandwidth, the instrument attenuated the peak and broadened the decay, yet still resolved the difference between the two currents, whereas reduced bandwidth or a longer sampling interval recovered only a fraction of the peak. The simulated trace serves as an instrument-independent reference.

### Procedure

### Electrode Fabrication on Glass Wafer**s**

#### Overview

Sensor chips are fabricated on Ti/Au-coated substrates by a two-level lithographic process. In the first level, a positive photoresist mask is used to define the metal electrode geometry, followed by sequential wet etching of Au and Ti. In the second level, SU-8 3005 is patterned to define the insulating/passivation layer while exposing the active sensing and contact regions. A final thin S1805 coating may be applied as a temporary protective layer for storage or downstream processing.

This process is suitable for rigid substrates and can be adapted to flexible substrates, although the hard-bake and cooling conditions must then be modified to minimize stress, cracking, or warping.

#### Reagents

- Cr quartz 5ʺ blank photomask with S1805 photoresist coating
- Ti/Au-coated substrates, prepared on glass (11774, platypustech)
- Acetone (CAS: 67-64-1)
- Isopropyl alcohol (IPA, 278475-Sigma-Aldrich)
- Deionized (DI) water
- S1805 positive photoresist (Fisher Scientific Catalog No.NC2003930)
- AZ 300 MIF developer (AZ® 300 MIF Photoresist Developer, EMD Performance Materials)
- AZ 1:4 400K developer (dilution with DI water; AZ® 400K Concentrate Photoresist Developer, EMD Performance Materials)
- Chromium etchant – (1020 Fisher Scientific Catalog No. NC9820871)
- Gold etchant - Type TFA (Fisher Scientific Catalog No. NC0977944)
- Titanium etchant – Type TFTN (Fisher Scientific No. NC0977946)
- SU-8 3005 negative photoresist (Kayaku Advanced Materials SU-8 3000)
- SU-8 developer (Fisher Scientific No. NC9901158)
- Nitrogen gas for drying

#### Reagent setup

*Ti/Au-coated substrates*.

Use substrates coated with titanium as an adhesion layer and gold as the electrode metal.

#### S1805

Bring to room temperature before use. Avoid introducing bubbles during dispensing. Do not use photoresist past its recommended shelf life.

#### SU-8 3005

Allow the bottle to equilibrate to room temperature before opening to avoid moisture condensation. Mix gently only if recommended by the manufacturer. Do not shake aggressively.

#### Wet etchants

Prepare and use according to institutional safety requirements and cleanroom wet-bench rules. Etchant performance depends strongly on age, temperature, and contamination.

#### Equipment

- Cleanroom or microfabrication workspace
- Spin coater
- Hotplates capable of 65 °C, 95 °C, 110 °C, 200 °C, and 135 °C if titanium etching is performed hot
- Mask aligner or UV exposure system
- Maskless aligner (Heidelberg MLA150)
- Plasma cleaner
- Nitrogen gun
- Chemical fume hood or wet bench
- Compatible wafer/substrate tweezers
- Optical microscope
- Profilometer or ellipsometer for resist thickness verification
- Timer
- Temperature-resistant chemical containers for etching

### Equipment setup

#### Mask aligner UV exposure system

Measure the exposure intensity at the wafer plane using a calibrated radiometer. The protocol below assumes 20 mW cm^-2^. If the real intensity differs, the exposure times must be adjusted accordingly.

The protocol below is recommended for 5ʺ chromium quartz masks pre-coated with S1805 photoresist.

**Table 1:**
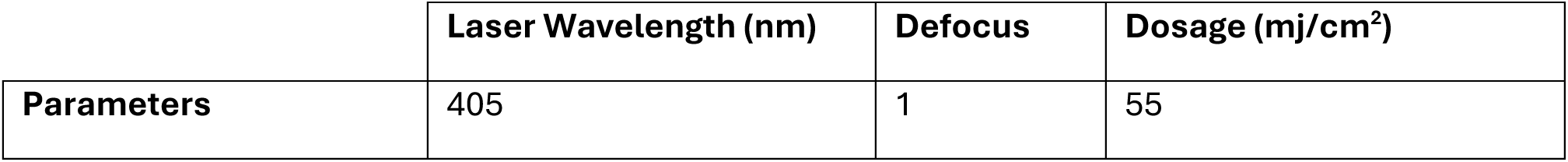
Parameters for mask aligner.

### Plasma cleaner

**Table 2:** Parameters for plasma cleaner.

|  | O<sub>2</sub> Flow (sccm) | Back Pressure (Pa) | Position (shelf/bottom) | RF Power (W) |
| --- | --- | --- | --- | --- |
| **Surface Treatment** | 20 | 20 | Shelf | 100 |
| **Descum** | 20 | 20 | bottom | 100 |

### Spin coater

**Table 3:** Parameters for spin coater.

| **Regent:** | **AZ nLOF 2035 & SurPass 4000** | **SU-8 3005** | **S1805** |
| --- | --- | --- | --- |
| Spread speed (rpm) | 500 | 500 | 500 |
| Spread acceleration (rpm/s) | 250 | 100 | 250 |
| Spread time (s) | 10 | 5 | 10 |
| Final speed (rpm) | 3500 | 4000 | 2000 |
| Final acceleration (rpm/s) | 1000 | 500 | 300 |
| Final spin time (s) | 30 | 30 | 40 |

#### Before beginning

All fabrication should be carried out using clean handling practices. Work in a cleanroom or low-particle environment. Handle substrates only with clean tweezers and powder-free gloves. Avoid touching coated or patterned surfaces. Minimize the time between cleaning, coating, exposure, and development steps.

The exact etch times and lithography conditions depend on substrate composition, metal thickness, resist age, humidity, lamp intensity, and mask quality. Before routine fabrication, calibrate each step using test substrates.

#### Procedure

## Part I. Fabrication of patterned Chromium masks

**●TIMING: 35-50 min, masks can be re-used for subsequent batches**

**1. Initial photomask cleaning (Timing:2-5 min)**

**(A) See Fig. 2a**

i. Blow dry thoroughly with nitrogen to remove any dust particles.
ii. Visually inspect mask for scratches on S1805 photoresist that might affect pattern.

**2. Mask exposure (Timing:20-30 min)**

i. Load mask into maskless aligner (Heidelberg MLA 150)
ii. Open the DXF design file, and specify the layer (Ti/Au pattern or SU-8 passivation)

a. This design requires two masks in total, one for each layer
b. Confirm that the dimensions are correctly to scaled
iii. Expose mask to UV radiation using the specified parameters for the maskless aligner

a. Laser wavelength: 405 nm, defocus: 1, dosage: 55mJ/cm^2^

**3. Mask photoresist development (Timing:2-3 min)**

i. Place mask into holder and submerge in AZ 1:4 400K for 1 min.
ii. Rinse with DI water.
iii. Dry thoroughly with nitrogen.

**4. Chromium etching (Timing:5-10 min)**

i. Place mask into holder and immerse in chromium etchant for 3 min, agitating gently.
ii. Rinse thoroughly with DI water.
iii. Dry with nitrogen.

*▲***Caution** Chromium etchants are corrosive and may be incompatible with some materials and tools.

**◆ Troubleshooting** If chrome residue remains in cleared regions, extend etching in 5–10 s increments until mask pattern is completely transparent.

## Part II. Patterning of Ti/Au electrodes using S1805

**●Timing: ∼45–75 min, excluding calibration**

**5. Initial substrate cleaning (Timing:5–10 min)**

i. Place the Ti/Au-coated substrate in a clean solvent-compatible holder.
ii. Sonicate in acetone for 2 min
iii. Sonicate in IPA for 2 min
iv. Sonicate in DI Water for 2 min
v. Blow dry for 2 min using nitrogen gun.

If using an automated solvent/developer station, the IPA and DI water steps may be performed in the tool, while acetone is added manually beforehand.

▲**Critical step** Residual organics, water, or particles will compromise resist adhesion and pattern quality.

*▲***Caution** Acetone and IPA are flammable. Use only in designated solvent handling areas.

**6. Coat S1805 positive photoresist (Timing: 3–5 min)**

i. Place the clean dry substrate on the spin coater chuck.
ii. Dispense sufficient S1805 to cover the substrate surface.
iii. Spin-coat at 2,000 rpm using the predefined recipe for S1805.

a. 500 rpm, acceleration 250 rpm s^-1^, 10 s
b. 2,000 rpm, acceleration 300 rpm s^-1^, 40 s

▲**Critical step** Avoid trapped bubbles and incomplete spreading. Nonuniform coating will produce variable development and poor etch definition.

▪ **Pause point** Coated substrates should proceed promptly to soft bake.

**7. Soft bake (Timing: 2–3 min)**

i. Bake the coated substrate on a hotplate at 110 °C for 70 s.
ii. Remove and allow to cool to room temperature for 2 min on a clean flat cleanroom wipe surface.

▲**Critical step** Do not overbake. Excess baking can reduce development rate and degrade feature transfer.

**8. UV exposure (Timing: 5–10 min)**

i. Align the substrate with the electrode photomask (layer ‘Ti/Au pattern’)
ii. Expose for 6 s at 20 mW cm^-2^. This corresponds to an approximate dose of 120 mJ cm^-2^.

▲**Critical step** Ensure good mask contact. Dust or mask gaps will blur the pattern edges.

**◆ Troubleshooting** If features consistently appear oversized or rounded, reduce exposure or improve mask contact. If development is incomplete, the exposure may be insufficient.

**9. Development of S1805 (Timing: 2–3 min)**

i. Develop manually in AZ 300 MIF for 1 min in a glass dish.
ii. Rinse immediately with DI water.
iii. Dry with nitrogen.
iv. Inspect under an optical microscope.

▲**Critical step** Do not let the substrate sit after development without rinsing.

*▲***Caution** AZ 300 MIF is corrosive and neurotoxic. Use in a ventilation hood, with nitrile gloves and appropriate eye protection.

**◆ Troubleshooting** If residue remains in cleared regions, extend development in 5–10 s increments. If resist lifting or severe edge erosion occurs, the resist may be underbaked or overdeveloped.

**10. Gold etching (Timing: 2–5 min)**

i. Immerse the developed substrate in gold etchant for approximately 70 s.
ii. Rinse thoroughly with DI water.
iii. Dry with nitrogen.

▲**Critical step** The correct Au etch time depends on gold thickness and etchant condition. Confirm complete Au removal on test pieces before processing valuable samples.

*▲***Caution** Gold etchants are corrosive and may be incompatible with some materials and tools.

**◆ Troubleshooting** If conductive remnants remain, Au was underetched. If the metal edge is badly recessed beneath the resist, the sample was overetched, or the etch is strongly isotropic.

**11. Titanium etching (Timing: 3–10 min)**

i. Etch the exposed Ti adhesion layer using preheated titanium etchant at 135 °C on a hot plate. For a 5–10 nm Ti layer, begin with a minimum etch time of 3 min. Inspect the sample visually. Continue etching in 30–60 s increments until the exposed Ti regions are completely removed. Do not exceed 5 min for 5–10 nm Ti unless incomplete removal is confirmed, as prolonged exposure may increase undercutting or damage adjacent metal features. The endpoint is reached when the exposed Ti has disappeared from the open regions, and no residual grey/dark Ti film is visible between or around the patterned Au features. Complete Ti removal can be confirmed by optical microscopy, loss of metallic contrast in the exposed regions, and, when available, profilometry, ellipsometry, sheet-resistance measurement, SEM/EDS or XPS on a test coupon processed in parallel. For new Ti thicknesses, new etchant lots, or different hot-plate conditions, first process a Ti-coated test coupon of the same thickness and determine the minimum time required for complete Ti removal. Use this value as the process endpoint for patterned devices.
ii. (ii) Rinse the sample immediately and thoroughly with DI water to stop the etch. Use a gentle but continuous rinse for at least 1 min, ensuring that no residual etchant remains on the surface or near patterned features.
iii. (iii) Dry the sample with nitrogen. Inspect the surface under an optical microscope to confirm complete removal of exposed Ti and to check for undercutting, delamination or damage to Au features.

**▲ Critical step** Monitor the etch closely. Do not leave the etch unattended.

**▲ Critical step** The etch rate depends strongly on Ti thickness, etchant formulation, bath age, temperature uniformity, sample loading and agitation. Do not rely only on nominal timing. Use endpoint inspection on the actual sample or on a matched test coupon.

*▲***Caution** Heated titanium etching is hazardous. Use only with proper wet-bench controls, splash protection, chemical-resistant gloves, eye protection, and approved high-temperature containers. Follow institutional safety rules.

## Part III. Cleaning and surface preparation before SU-8 Timing: ∼20 min

**12. Solvent rinse after metal etch**

i. Rinse the patterned substrate in acetone for 2 min.
ii. Transfer to IPA for 2 min.
iii. Rinse with DI water.
iv. Dry thoroughly with nitrogen.

▲**Critical step** Ensure that all etchant and S1805 residues are completely removed. Residual acid or salts can compromise SU-8 adhesion. Use a voltmeter to check for unexpected shorts caused by residual titanium.

**13. Plasma cleaning (Timing: ∼5 min)**

i. Place the substrate in the plasma cleaner.
ii. Plasma-clean for 3 min with specified parameters for surface treatment.

a. O_2_ flow: 20 sccm, back pressure: 20 Pa, sample position: shelf, RF power: 100 W

▴**Critical step** Use the minimum plasma dose needed for cleaning and activation. Overexposure can damage surfaces or alter surface chemistry.

**14. Dehydration bake (Timing: ∼12 min)**

i. Transfer the plasma-cleaned substrate to a hotplate.
ii. Bake at 200 °C for 10 min.

This step acts as a dehydration bake and can improve subsequent SU-8 coating and adhesion.

▲**Critical step** After dehydration, avoid prolonged exposure to humid air before SU-8 coating.

**Part IV. SU-8 passivation patterning Timing: ∼45–75 min**

**15. Coat SU-8 3005 (Timing: 3–5 min)**

i. Place the substrate on the spin coater chuck.
ii. Dispense SU-8 3005 onto the center of the substrate.
iii. Spin-coat using the 4,000 rpm recipe.

a. 500 rpm, acceleration 100 rpm s^-1^, 5 s
b. 4,000 rpm, acceleration 500 rpm s^-1^, 30 s

**16. SU-8 soft bake (Timing: 6–8 min)**

i. Bake the coated substrate at 95 °C for 5 min.
ii. Remove and allow the substrate to cool briefly.

▲**Critical step** Avoid rapid temperature shock. Internal stress in SU-8 can cause cracking or adhesion failure later.

**17. SU-8 exposure (Timing: 5–10 min)**

i. Align the substrate to the passivation photomask (layer ‘SU-8 passivation’).
ii. Expose for 15 s at 20 mW cm^-2^. This corresponds to an approximate dose of 300 mJ cm^-2^.

▲**Critical step** Alignment is crucial because this layer determines the exposed sensing and contact windows.

**◆ Troubleshooting** If passivation openings shrink or close after development, overexposure is likely. If SU-8 lifts off or remains soft, exposure or post-exposure bake may be insufficient.

**18. Post-exposure bake (Timing: 5–7 min)**

i. Bake at 65 °C for 1 min.
ii. Increase to 95 °C and bake for 3 min.

▲**Critical step** This step completes crosslinking. Poor control here causes incomplete development, weak features, or residual film in open windows.

**19. Development of SU-8 (Timing: 3–8 min)**

i. Develop the substrate in SU-8 developer for 50 s.
ii. Remove the substrate from developer solution and rinse with IPA for 10 s.
iii. Inspect visually.
iv. If a white cloudy film appears, return the substrate to SU-8 developer for 5 s.
v. Rinse again with IPA.
vi. Repeat as needed until no cloudy residue remains.
vii. Dry with nitrogen.

▲**Critical step** Cloudiness after IPA is a sign of incomplete development. Do not ignore it.

**◆ Troubleshooting** Persistent cloudiness may indicate underdevelopment, underexposure, poor post-exposure bake, or excessively thick SU-8.

**20. Hard bake (Timing: ∼45–60 min including ramping)**

i. Place the developed substrate on a hotplate.
ii. Ramp the temperature gradually to 200 °C (Start at 65°C and ramp to 200C (as long as it takes))
iii. Hold at 200 °C for 30 min.
iv. Ramp down to 65°C gradually and then room temperature.

For flexible substrates, use slower heating and cooling ramps to minimize warping or film cracking.

▲**Critical step** Do not place thin flexible substrates directly onto a fully equilibrated 200 °C hotplate without ramping.

*▲***Caution** Confirm that the substrate, metal stack, and any supporting adhesive or carrier can tolerate 200 °C.

## Part V. Temporary protective photoresist coating Timing: ∼10 min

**21. Apply protective S1805 layer**

i. Spin-coat S1805 at 1,000 rpm.
ii. Bake at 110 °C for 60 s.

This protective coating may be used to shield the device surface during handling, storage, dicing, or further processing.

▲**Critical step** If later removal is required, verify compatibility with the SU-8 layer and exposed sensing interfaces.

Timing

Initial cleaning: 5–10 min
S1805 coating and bake: 5–8 min
Exposure and development: 7–12 min
Au and Ti etching: 5–15 min
Post-etch cleaning and plasma/dehydration: 15–20 min
SU-8 coating and bake: 8–12 min
SU-8 exposure and post-exposure bake: 8–12 min
SU-8 development: 3–8 min
Hard bake with ramping/cooling: 45–60 min
Protective resist coating: 8–10 min

Total: ∼2.0–3.5 h per batch, excluding mask fabrication, mask alignment optimization and etch calibration

### ▴ Critical steps

**(i) Substrate cleanliness:** Dirty substrates cause adhesion failure, pinholes, and pattern collapse.
**(ii) Exact etch calibration:** Au and Ti etch times are batch dependent. Never trust nominal times blindly.
**(iii) Humidity control before SU-8:** Moisture kills SU-8 consistency and adhesion.
**(iv) True thickness measurement:** Measure S1805 and SU-8 thickness to ensure the correct dimensions.
**(v) Careful Ti etching:** This is the most hazardous and variable step.
**(vi) Cloudiness after SU-8 IPA rinse:** This means incomplete development. Do not proceed until resolved.
**(vii) Controlled hard-bake ramps:** Especially critical for flexible substrates.

**◆ Troubleshooting** See Table 4.

**Table 4:** Troubleshooting the electrode fabrication.

| Step | Problem | Possible reason | Solution |
| --- | --- | --- | --- |
| S1805 coating | Streaks or nonuniform film | Dirty substrate, poor spread, resist bubbles | Improve cleaning, allow resist to spread, degas resist if needed |
| S1805 development | Residue remains in open regions | Underdevelopment or insufficient exposure | Extend development slightly or verify exposure intensity |
| Metal etch | Residual conductive paths | Underetching | Increase etch time in small increments after calibration |
| Metal etch | Severe edge undercut | Overetching or isotropic etch | Reduce etch time, improve control of etchant condition |
| SU-8 coating | Cracks after bake | Thermal stress or excessive dehydration | Use gradual ramps and verify film thickness |
| SU-8 development | White cloudiness after IPA | Incomplete development | Return briefly to developer and repeat IPA rinse |
| SU-8 pattern | Openings smaller than mask | Overexposure or excessive post-exposure bake | Reduce dose or post-exposure bake |
| SU-8 delamination | Poor adhesion | Moisture, residue, insufficient dehydration | Improve post-etch cleaning and dehydration bake |

### Anticipated results

**When the process is properly optimized, the fabricated chips should show:**

well-defined Ti/Au pattern, similar to the designed mask, with clean edges,
complete electrical isolation of passivated regions,
accurately opened SU-8 windows over the sensing and contact pads,
no residual metal bridging between features,
no visible SU-8 residue in exposed windows,
no major cracking or delamination of the passivation layer.

Under an optical microscope, metal features should exhibit uniform edge definition, and passivation openings should align cleanly with the underlying electrode structures. If profilometry is available, the SU-8 thickness and step heights should match the values reported in the fabrication specifications.

### ***▴***Cautions and safety notes

**Solvents** Acetone and IPA are flammable and volatile. Use appropriate ventilation and ignition-free handling.

**UV exposure** Avoid direct eye and skin exposure to UV radiation. Use the aligner according to facility procedures.

**Gold etchant** Corrosive. Can damage nearby materials and equipment if mishandled.

**AZ 300 MIF** Corrosive and contains TMAH, a severe neurotoxin. Use appropriate ventilation and PPE.

**Hot titanium etch** Includes heating corrosive chemistry and must be performed in a fume hood. Use proper PPE and include facility-specific handling requirements.

**Plasma cleaner** Can modify surfaces aggressively if overused. Do not overexpose delicate structures.

**SU-8** Uncured SU-8 and developer require careful chemical handling. Avoid skin contact and cross-contamination.

**High-temperature hard bake** Thermal mismatch between substrate, metal, and polymer layers can cause cracking, warping, or delamination.

**Ti/Au glass fabrication** In some experiments, patterned Ti/Au glass wafers were prepared in-house rather than sourced commercially. In these cases, glass wafers were first patterned with a lift-off resist, metallized by electron-beam deposition of Ti/Au, and then subjected to solvent lift-off before downstream device fabrication.

[Optional] Upstream procedure: preparation of Ti/Au-patterned glass wafers Reagents

Glass wafers
Acetone
Isopropyl alcohol (IPA)
Deionized (DI) water
AZ nLOF 2035 negative liftoff photoresist (AZnLOF2035-Q, AZ® nLoF 2035 Negative Tone Lift-Off Photoresist, EMD Performance Materials)
SurPass 4000 adhesion promotor
AZ 300 MIF developer or equivalent alkaline developer (AZ300MIF-CS, AZ® 300 MIF Photoresist Developer, EMD Performance Materials)
Titanium source for electron-beam evaporation
Gold source for electron-beam evaporation
Nitrogen gas for drying

### Equipment

Cleanroom or microfabrication workspace
Spin coater
Hotplate set to 110 °C
Mask aligner (MA6)
Plasma cleaner (Samco 300)
Electron-beam evaporator (AJA)
Solvent-compatible wafer holder or container
Solvent-compatible tweezers for wafer handling
Nitrogen gun
Pipette for gentle solvent-assisted lift-off
Optical microscope

**22. Cleaning of glass wafers (Timing: 5–10 min)**

i. Rinse the glass wafer sequentially with acetone, IPA, and DI water.
ii. Dry thoroughly using a nitrogen gun.

▲**Critical step** The wafer surface must be free of organic residue, particles, and water before resist coating. Surface contamination will reduce adhesion and can cause incomplete lift-off or metal defects.

*▲***Caution** Acetone and IPA are flammable and should be handled in an approved solvent-processing area.

**23. Coating with adhesion promoter (Timing: 3–5 min)**

i. Place the clean glass wafer on the spin coater chuck.
ii. Dispense SurPass 4000 adhesion promoter onto the wafer surface.
iii. Spin-coat using the following program:
iv. 500 rpm, acceleration 250 rpm s^-1^, 10 s
v. 3,500 rpm, acceleration 1,000 rpm s^-1^, 30 s

▴**Critical step** Ensure complete wafer coverage during dispense. Incomplete coverage can lead to nonuniform photoresist adhesion and pattern failure during development or lift-off.

**24. Coating with nLOF 2035 photoresist (Timing: 3–5 min)**

i. Dispense nLOF 2035 photoresist onto the wafer.
ii. Spin-coat using the same program:
iii. 500 rpm, acceleration 250 rpm s^-1^, 10 s
iv. 3,500 rpm, acceleration 1,000 rpm s^-1^, 30 s

▲**Critical step** Avoid bubbles and resist streaking. Nonuniform resist thickness will affect exposure, development, and the quality of the lift-off profile.

**25. Soft bake (Timing: ∼2 min)**

i. Bake the coated wafer on a hotplate at 110 °C for 1 min.
ii. Allow the wafer to cool briefly on a clean flat surface.

▲**Critical step** Do not overbake. Excess baking may reduce the resist contrast and compromise clean lift-off.

**26. UV exposure (Timing: 5–10 min)**

i. Align the wafer with the photomask.
ii. Ensure that the wafer is centered properly before exposure.
iii. Expose for 5 s at 20 mW cm^-2^. This corresponds to an approximate dose of 100 mJ cm^-2^The same mask for patterning S1805 on Platypus wafers may be used here.

*▲***Critical step** Correct wafer centering and mask alignment are essential for uniform feature placement across the wafer.

**27. Post-exposure bake (Timing: ∼2–3 min)**

i. Bake the exposed wafer at 110 °C for 90 s.
ii. Allow the wafer to cool briefly.

▲**Critical step** This post-exposure bake should be consistent across batches, as it affects feature definition and development behavior.

**28. Development (Timing: 3–5 min)**

i. Develop the wafer in AZ 300 MIF for 1 min.
ii. Rinse thoroughly with DI water.
iii. Dry with nitrogen.
iv. Inspect the developed pattern under an optical microscope.

▲**Critical step** Incomplete development will cause poor lift-off and unwanted metal bridging. Overdevelopment can distort feature dimensions.

*▲***Caution** AZ 300 MIF is corrosive and neurotoxic. Use in a ventilation hood, with nitrile gloves and appropriate eye protection.

**◆ Troubleshooting** If residual resist remains in exposed areas, extend development slightly in 5–10 s increments. If features are badly eroded, reduce development time or verify exposure conditions.

**29. Plasma cleaning (Timing: ∼5 min)**

i. Plasma-clean the developed wafer for 2 min using the defined descum recipe.
ii. O_2_ flow: 20 sccm, back pressure: 20 Pa, sample position: bottom, RF power: 100 W

▲**Critical step** This step should be used consistently and only as needed. Excessive plasma treatment can alter resist sidewalls and affect lift-off performance.

**30. Electron-beam evaporation of Ti/Au (Timing: ∼1.5–2.5 h including pump-down)**

i. Load the patterned wafers into the electron-beam evaporator.
ii. Pump down the chamber for approximately 1 h, until the pressure falls below 2 × 10^-6^ Torr.
iii. (iii) Deposit a 10 nm titanium adhesion layer at a rate of 0.03 nm/s.
iv. (iv) Without breaking vacuum, deposit 100 nm gold at a rate of 0.05 nm/s.
v. (v) Vent the chamber.
vi. (vi) Remove the wafers.

▲**Critical step** Do not vent before the target base pressure is reached. Poor vacuum will degrade film quality, adhesion, and reproducibility.

▲**Critical step** Titanium should be deposited before gold without breaking vacuum. Otherwise, adhesion and interface quality may suffer.

**31. Lift-off of excess metal (Timing: ≥1 h)**

i. Immediately place the metallized wafer into a covered acetone bath.
ii. Leave the wafer submerged for at least 1 h.
iii. While keeping the wafer immersed in acetone, use a pipette to gently direct solvent flow across the surface and assist removal of excess metal.
iv. Continue until all unwanted metal has lifted off and only the intended patterned metal remains.
v. Once lift-off is complete, rinse the wafer sequentially with acetone, IPA, and DI water.
vi. Dry with nitrogen.

▲**Critical step** The wafer must remain fully covered in acetone until all excess metal has been removed. Once the wafer dries, or once it is exposed to DI water before lift-off is complete, the residual metal flakes can redeposit or become extremely difficult to remove.

*▲***Major caution** Do not allow the wafer to dry and do not expose it to DI water before lift-off is complete.

**◆ Troubleshooting** If excess gold does not lift off cleanly, likely causes include inadequate resist profile, insufficient development, excessive metal sidewall coverage, or premature drying during lift-off.

*▴***Caution** Do not use aggressive mechanical scraping. That will destroy fine features or delaminate the patterned metal.

**Timing**

Wafer cleaning: 5–10 min
Adhesion promoter coating: 3–5 min
nLOF 2035 coating: 3–5 min
Soft bake: 2–3 min
Exposure and alignment: 5–10 min
Post-exposure bake: 2–3 min
Development: 3–5 min
Plasma clean: 5 min
Electron-beam loading and pump-down: ∼1 h
Ti/Au deposition: 20–45 min, depending on deposition rates
Lift-off: at least 1 h

Total: ∼3–4.5 h, excluding queue time for the evaporator

### ▴ Critical steps

**Wafer cleanliness before resist coating:** Any residue will compromise adhesion and lift-off.

**Uniform SurPass 4000 and nLOF 2035 coating:** Poor uniformity causes variable feature quality.

**Correct centering before exposure:** Miscentering leads to mask-placement errors across the wafer.

**Complete development before metal deposition:** Residual resist leads to metal bridging and failed lift-off.

**Adequate vacuum before deposition:** Do not deposit at poor base pressure.

**No premature drying during lift-off:** This is the most important practical caution in this process.

### Alternative silicon-based fabrication routes

Although the procedure described above focuses on fabrication of the sensor chip on glass substrates, the same electrode architecture can also be implemented on silicon wafers using a related microfabrication workflow and an alternative insulating layer to define the

∼20 µm sensing aperture. This silicon-based route may be useful when integration with other microfabricated components or substrate-specific processing steps is desired. A brief overview of the silicon-based fabrication procedure and the corresponding aperture-definition strategy is provided in Box 3^44,45^.

#### BOX 3 Other cleanroom processes and devices

Cleanroom fabrication facilities are among the most expensive laboratory environments required for device development. While commercial semiconductor foundries provide access to such infrastructure, many universities and research institutes also maintain cleanroom facilities supported by government funding and industry partnerships to enable microfabrication-based research and development. In this protocol, SU8-based processing is the primary approach for fabricating miniaturized devices compatible with molecular pendulum technology. However, for laboratories with established silicon microfabrication capabilities, an alternative route based on silicon/silicon nitride processing can also be used to design and manufacture miniaturized chips for electrochemical interfaces.

The device manufacturing process progresses through process development within each cleanroom facility, and the specific fabrication recipes may vary depending on the available equipment and infrastructure. Therefore, the fabrication steps should be carefully optimized during process development to ensure reliable device production.

Example process of Si/Au/Si_3_N_4_ devices: This process requires two masks: a metal mask and a passivation mask.

**Silicon processing:** An oxide layer (Silicon dioxide, SiO_2_) is thermally grown on the silicon wafer. The thickness of the oxide may vary from 300 nm to a few microns.

**Lithography:** A metal mask is used to transfer the designed pattern onto the substrate. The wafer is first treated with hexamethyldisilazane (HMDS) to promote adhesion, followed by spin coating of LOR3A. The exposed resist is developed in MF-319 developer, rinsed with deionized (DI) water, and dried under a nitrogen (N₂) stream.

**Lift-off:** The wafers are loaded into the deposition chamber for argon plasma treatment, followed by metal deposition. Titanium and gold layers are deposited by evaporation (Ti, 10–20 nm; Au, 150 nm). Excess metal is removed by lift-off process by immersing the wafer in R1165 resist remover until the unwanted metal is released. The wafer is then rinsed thoroughly with DI water and dried under a N₂ stream.

**Note:** After the metal deposition, apply a piece of cleanroom tape (Kapton®) onto the metal surface and press gently. Peel the tape off. The tape test allows evaluation of the metal adhesion on the substrate. After lift-off, check the wafer under the microscope.

Passivation and Etch: Silicon nitride is deposited onto the substrate with a thickness of 300–1000 nm to serve as a passivation layer. S1813 photoresist is then spin-coated and processed according to standard lithography procedures. The wafer is subsequently exposed using the passivation mask and developed in MF-319 developer. After development, the wafer is rinsed with deionized (DI) water and dried under a nitrogen (N₂) stream. The exposed silicon nitride is then etched using inductively coupled plasma (ICP) etching. Following the etching step, the remaining photoresist is removed by immersing the wafer in R1165 resist remover. The wafer is finally rinsed thoroughly with DI water and dried under a nitrogen (N₂) stream.

**Note:** Check the wafer under the microscope, Atomic Force Microscopy (AFM) and/or ellipsometer. The passivation thickness should be decided based on the application since it may play a critical role in the electrochemistry.

**Dicing:** The wafer is coated with S1813 photoresist as a protective layer prior to dicing. Individual devices are subsequently obtained by mechanical wafer dicing.

**Note:** Wafer-scale fabrication yields multiple devices on a single wafer. These wafers should be stored at room temperature in sealed wafer holders to minimize contamination. Each device should be washed with organic solvents such as acetone and isopropyl alcohol before their use. Another alternative is the use of a resist remover.

**Fig. 1.**
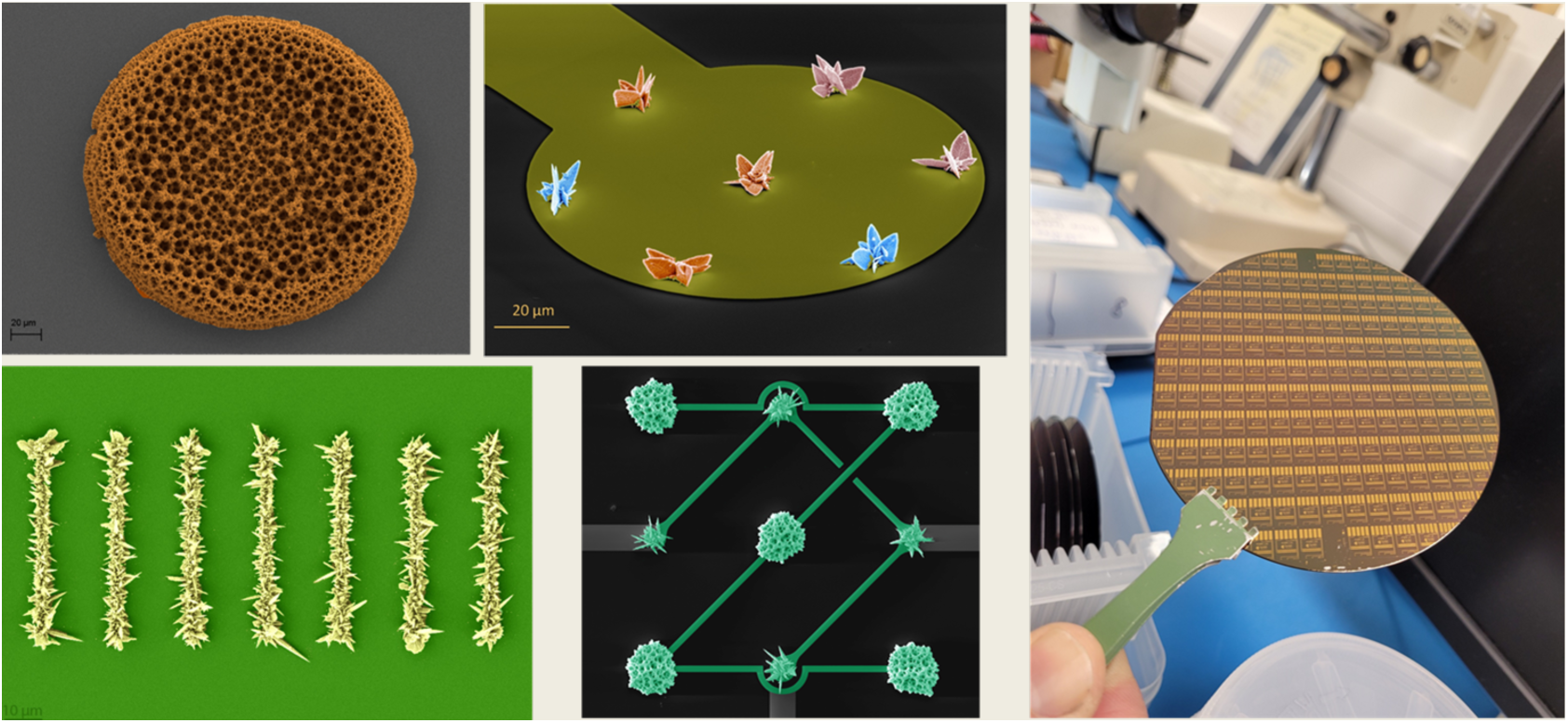
Representative silicon-based chips on a wafer and SEM images of nanostructured silicon-based devices.

### Biolayer interferometry (BLI) for affinity determination

#### (A) Experimental design

Before the development of the electrochemical sensors for protein measurement, it is critical to characterize affinity reagents used for sensor development, given the variability of both commercial reagents and those reported in the scientific literature. BLI can be used to determine the binding affinity of recognition elements, including antibodies, aptamers, and nanobodies, toward their corresponding targets under label-free conditions by monitoring changes in optical thickness at the biosensor surface during association and dissociation (Box 4). It is strongly recommended that all affinity reagents be validated with BLI before constructing or testing pendulum sensors. In a typical experiment, the recognition element is immobilized on a biosensor compatible with its chemistry, a stable baseline is established in the assay buffer, the target is introduced at a series of concentrations to monitor association, and the sensor is then returned to the buffer to measure dissociation. The resulting sensorgrams are analyzed using either kinetic or steady-state fitting to obtain an apparent equilibrium dissociation constant (K_D_). Because the apparent affinity can depend strongly on buffer composition, surface loading density, nonspecific adsorption, and the selected fitting model, these parameters should be optimized for each receptor–target pair rather than treated as universal. It is also important to note that BLI measurements are typically performed in assay buffers containing blocking additives such as bovine serum albumin (BSA) and Tween 20 to reduce nonspecific adsorption, whereas MP sensors rely on an electrochemical interface passivated with surface-bound molecules such as 6-Mercapto-1-hexanol (MCH). As a result, the K_D_ obtained by BLI should be regarded as an apparent affinity measured under BLI-specific interfacial conditions and may not fully capture the binding kinetics governing MP sensing at the electrode surface.

For antibody-based measurements, capture biosensors compatible with the antibody format can be used. For aptamer-based measurements, biotinylated aptamers can be immobilized on streptavidin biosensors. For nanobody-based measurements, VHH-compatible biosensors can be used. In all cases, the immobilization format should preserve receptor activity and minimize steric or orientational constraints that could distort the measured binding response.

Reagents

Recognition element of interest, such as antibody, aptamer, or nanobody
Target analyte, prepared over an appropriate concentration range
Assay buffer, optimized for the receptor–target system
Optional buffer additives to reduce nonspecific adsorption and improve baseline stability, including BSA and Tween 20
Biosensors appropriate for the immobilization strategy, such as capture biosensors, streptavidin biosensors, or VHH-compatible biosensors
Ultrapure water
Low-binding sample tubes or BLI-compatible microplates

Reagent setup Assay buffer

The assay buffer should be optimized for each receptor–target system. Depending on the biological application and the properties of the binding pair, suitable buffers may include phosphate buffered saline (PBS) alone or PBS supplemented with BSA and/or Tween 20 to reduce nonspecific adsorption and improve signal stability. Buffer composition, pH, ionic strength, and surfactant content can all affect the apparent affinity and should therefore be selected carefully.

Target concentration series

Prepare a target dilution series spanning below, near, and above the expected K_D_. The exact concentration window should be adjusted for each system on the basis of prior knowledge or exploratory measurements.

Equipment

Biolayer interferometry instrument (Octet® BLI, Sartorius)
Biosensor tips compatible with the receptor immobilization method
BLI-compatible microplate
Agitation-capable measurement system
Temperature-controlled measurement chamber, if available
Pipettes and low-retention pipette tips
Data-acquisition and analysis software

#### **(B)** Procedure

**32. Select the biosensor chemistry (Timing: 5–10 min)**

i. Select a biosensor compatible with the recognition element and immobilization strategy.
ii. For aptamer measurements, use streptavidin biosensors with biotinylated aptamers.
iii. For nanobody measurements, use VHH-compatible biosensors.
iv. For antibody measurements, use a capture biosensor suitable for the antibody format.

▴**Critical step** The immobilization strategy should preserve receptor accessibility and minimize steric hindrance. Poor orientation or surface crowding can distort both kinetic behavior and the apparent K_D_.

**33. Prepare assay buffer and samples (Timing: 15–30 min)**

i. Prepare the assay buffer using a formulation suitable for the receptor–target system.
ii. Dispense assay buffer, receptor solution, target concentration series, and control solutions into the BLI plate.
iii. Include buffer-only wells for baseline and dissociation measurements.
iv. Prepare control wells to evaluate nonspecific binding and baseline quality.

▲**Critical step** The buffer should be optimized for each system. Depending on the receptor and target, suitable formulations may include PBS alone or PBS supplemented with BSA and/or Tween

20. If the baseline is unstable or the control response resembles the main binding response, the buffer should be re-optimized before proceeding.

**34. Hydrate biosensors (Timing: 10–20 min)**

i. Hydrate the biosensors in the assay buffer for 10 min, or according to the manufacturer’s recommendation.

▲**Critical step** Incomplete hydration can lead to unstable baselines and poor reproducibility.

**35. Establish the initial baseline (Timing: variable, typically 60–300 s or longer)**

i. Transfer hydrated biosensors into the assay buffer and record the initial baseline.

▲**Critical step** Baseline duration is system-dependent and should not be fixed arbitrarily. Some receptor–sensor combinations stabilize rapidly, whereas others require longer equilibration. Establishing a stable baseline is essential for reliable affinity determination.

*▲***Caution** Do not proceed to loading or association if the baseline continues to drift significantly.

**36. Load or capture the receptor (Timing: variable)**

i. Transfer the biosensors into the receptor solution and monitor receptor loading in real time.
ii. Continue loading until a suitable and stable response is obtained for the selected system.

Because high surface loading can bias the measured binding response through mass-transport limitation, receptor density should be optimized for each system rather than maximized indiscriminately (Box 4).

▲**Critical step** Receptor loading density should be optimized for each biorecognition element separately, as suggested in the manufacturer’s manuals for different types of BLI biosensors. Excessive loading can cause steric crowding, rebinding, or mass-transport effects, whereas insufficient loading can reduce signal-to-noise ratio.

*▲***Caution** Higher loading does not necessarily improve the measurement. Overloading frequently degrades data quality.

**37. Re-establish the post-loading baseline (Timing: variable, typically 60–300 s)**

i. Transfer the loaded biosensors into the assay buffer to remove loosely bound material and re-establish a stable post-loading baseline.

▲**Critical step** This step is important for separating true target binding from drift caused by incomplete stabilization of the immobilized receptor.

**38. Measure association (Timing: variable)**

i. Transfer the loaded biosensors into wells containing the target at a series of concentrations. Note: suggested ranges are in the manufacturer’s manual.
ii. Record the association phase for a duration appropriate to the interaction kinetics (10 to 15 min).

▲**Critical step** Association time should be selected empirically. Fast interactions may be adequately captured in short association windows, whereas slower systems may require substantially longer measurements.

**39. Measure dissociation (Timing: variable)**

i. Transfer the biosensors from the target wells back into the assay buffer and record the dissociation phase (10 to 15 min).

▴**Critical step** Dissociation should be monitored long enough to resolve meaningful decay. Very slow dissociation may require extended measurement times.

**40. Include control conditions (Timing: throughout the experiment)**

i. Include a sensor without a receptor exposed to the target protein to assess nonspecific adsorption to the sensor surface.
ii. Include receptor-loaded sensors exposed only to the buffer to assess baseline drift in the absence of a target.
iii. Include receptor-loaded sensors exposed to a nontarget protein to confirm minimal off-target response.
iv. Subtract the appropriate control signal from the main binding curves during analysis.

▲**Critical step** If the control curve resembles the main binding curve, or if the baseline is unstable, the assay conditions are not yet optimized and the buffer composition should be adjusted before affinity values are interpreted.

#### **(C)** Data analysis

1. **Preprocess sensorgrams (Timing: 15–60 min)**
2. Inspect all sensorgrams for baseline stability, loading consistency, and concentration-dependent response.
3. Apply reference subtraction using the appropriate control traces.
4. Exclude traces showing severe drift, poor loading, or obvious artifacts.

▲**Critical step** Reference subtraction is essential when nonspecific binding or baseline drift is present.

**42. Fit the binding response (Timing: 15–60 min)**

i. Fit the association and dissociation curves using an appropriate binding model in the BLI analysis software.
ii. Extract the apparent K_D_ using either global fitting across concentrations or individual fitting of each concentration trace, depending on data quality.
iii. If individual fits are used, calculate an average K_D_ across the accepted traces.

▲**Critical step** Global fitting and individual fitting can yield different K_D_ values. In some cases, global fitting may not adequately describe both the association and dissociation phases, particularly when the dataset deviates from the assumptions of the fitting model. Under such conditions, fitting each concentration trace individually and averaging the resulting K_D_ values may provide a more representative estimate.

▲**Critical step** The apparent K_D_ is model dependent. Different preprocessing choices, fitting strategies, and trace-selection criteria can alter the reported affinity. These choices should therefore be reported explicitly.

*▲***Caution** A mathematically acceptable fit is not necessarily physically meaningful. Residuals, concentration dependence, and consistency across replicates should all be evaluated before accepting the final K_D_.

Timing

Sample and buffer preparation: 15–30 min
Biosensor hydration: 10–20 min
Initial baseline: variable
Receptor loading: variable
Post-loading baseline: variable
Association: variable
Dissociation: variable
Data analysis: 30–120 min

### Anticipated results

A successful experiment should produce stable baselines, reproducible receptor loading, concentration-dependent association responses, and dissociation curves suitable for fitting. Under optimized conditions, the extracted apparent K_D_ should be reasonably consistent across replicate measurements and fitting approaches. Large discrepancies between global and individual fits, substantial control responses, or unstable baselines indicate that assay conditions or model assumptions require further optimization.

- **◆** Troubleshooting

**Problem:** Baseline drift before association

**Possible reason:** Incomplete hydration, unstable receptor immobilization, or suboptimal buffer

**Solution:** Increase hydration time, extend baseline equilibration, and optimize buffer composition

**Problem:** Control trace resembles the main trace

**Possible reason:** Nonspecific adsorption or poor buffer conditions

**Solution:** Re-optimize the buffer, including possible use of PBS, BSA, and/or Tween 20 at optimal percentages, and verify surface blocking conditions

**Problem:** Weak binding signal

**Possible reason:** Insufficient receptor loading, low target concentration, or poor receptor activity

**Solution:** Optimize receptor loading concentrations, verify receptor functionality, and expand the concentration range of the target

**Problem:** Global fit does not describe the curves well

**Possible reason:** Model mismatch, heterogeneous surface behavior, or nonideal kinetics

**Solution:** Fit individual concentration traces, compare the resulting K_D_ values, and report the fitting strategy clearly

**Problem:** Apparent affinity changes with receptor loading

**Possible reason:** Surface crowding, rebinding, or mass-transport limitation

**Solution:** Reduce loading density and reassess the fit under less crowded conditions

#### BOX 4 Validation of affinity receptors by BLI

The concentration range used for target incubation should be selected in relation to the binding affinity of the receptor, typically expressed as the apparent dissociation constant (K_D_). In general – while pendulum sensors often show responses at low concentration levels – initial sensor development efforts are most informative when tested across concentrations spanning below, near, and above the K_D_, as this range captures the transition from low occupancy to near-saturation of the sensing interface. Because different recognition elements exhibit substantially different affinities, the optimal calibration window is receptor-dependent. In addition, the apparent affinity can be influenced by experimental conditions, including biofluid composition, pH, ionic strength, and temperature. Representative examples are shown here for TNF-α detected using a nanobody and for IL-1β detected using an antibody (Fig. 1). K_D_ values under a single set of conditions were determined by biolayer interferometry (BLI). It is noteworthy that K_D_ values are strongly affected by solution conditions and by the sensitivity of the instrumentation used.

**Fig. 1.**
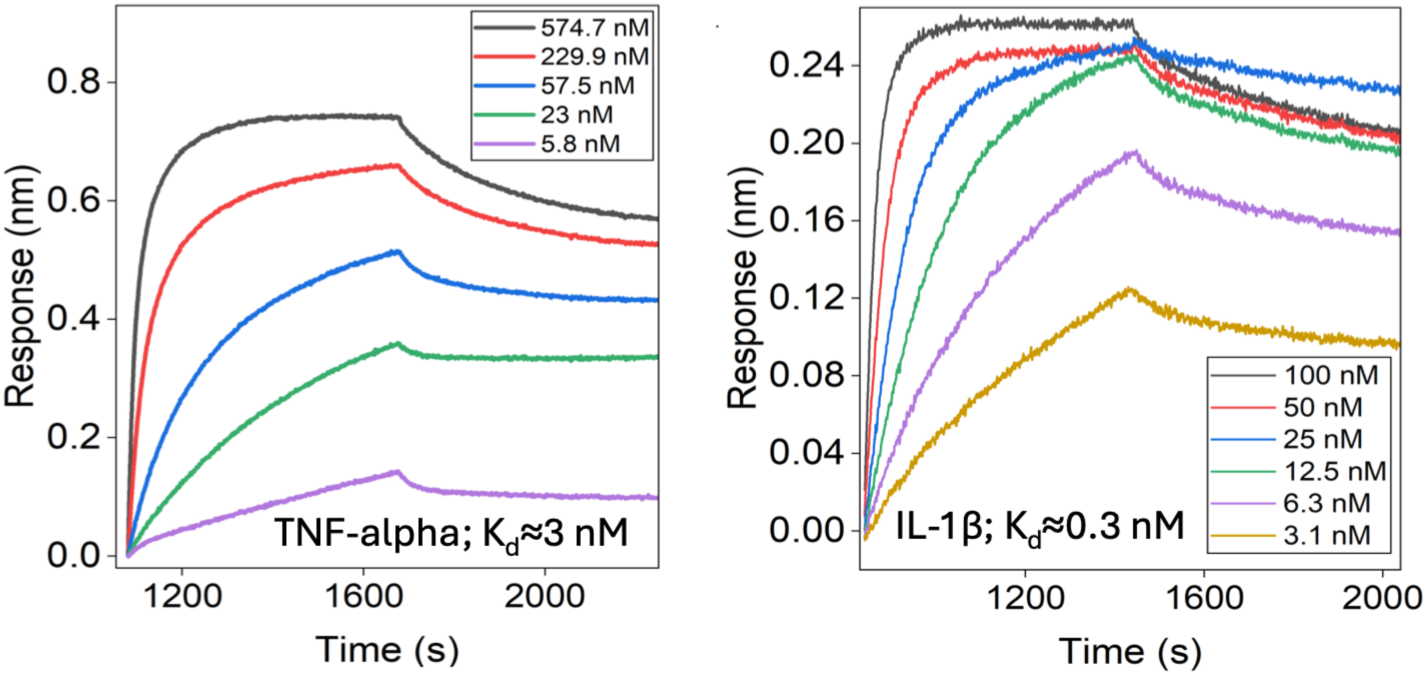
BLI assessment of TNF-α detected using a nanobody, and IL-1β detected using an antibody.

##### Mass-transport limitation in biolayer interferometry

Mass-transport limitation occurs when delivery of the target from bulk solution to the biosensor surface is slower than the intrinsic binding reaction between the target and the immobilized receptor. Under these conditions, the measured association phase is influenced not only by the true molecular interaction but also by diffusion and mixing, such that the recorded sensorgram no longer reflects purely reaction-limited binding. As a result, the fitted kinetic constants may be distorted, and the apparent K_D_ may deviate from the intrinsic affinity of the receptor–target pair.

This effect is more likely to occur when receptor loading is too high, the interaction is very fast, the target concentration is low, or agitation is insufficient. In practice, mass-transport limitation may be suspected if association curves change disproportionately with receptor loading, if global fits fail to capture the early association phase, or if fitted kinetic parameters vary strongly with sensor loading density. To reduce this effect, receptor loading should be lowered, agitation should be maximized within instrument limits, and fitting results should be compared across different loading conditions. When such effects are present, the measured K_D_ should be interpreted as an apparent surface-dependent value rather than a purely intrinsic binding constant.

##### Take-home message

Receptor loading in BLI should be optimized not only to maximize signal but also to avoid transport-limited binding, which can artificially distort both kinetic parameters and affinity estimates.

### Sensor preparation and target measurement

#### **(A)** Experimental Design

The MP-based assay utilizes a DNA-based tether system. A thiolated probe (P1) is immobilized onto a gold NME, while a secondary probe (P2), conjugated to either an antibody, a nanobody or an aptamer, hybridizes to form the pendulum structure. Binding of the target biomarker alters the electrochemical dynamics of the ferrocene redox reporter, as monitored by CA.

### Reagents

**Patterned sensor chips:** glass chips bearing patterned Ti/Au electrodes (5 nm Ti adhesion layer, 100 nm Au layer) with a 20 µm aperture defining the active sensing area.
**Gold precursor solution:** hydrogen tetrachloroaurate(III) hydrate (HAuCl_4_; 484385, Sigma-Aldrich) for electrodeposition of nanostructured microelectrodes (NMEs).
**Hydrochloric acid (HCl; 7647-01-0, Fisher Scientific)** for preparation of the gold electrodeposition solution.
**Sulfuric acid (H_2_SO_4_; 7664-G3-G, Fisher Scientific),** 100 mM, for electrochemical quality-control measurements by cyclic voltammetry (CV).
**Acetone** for chip cleaning.
**Isopropyl alcohol (IPA)** for chip cleaning.
**Deionized (DI) water** for reagent preparation and rinsing steps.
**Phosphate-buffered saline (PBS; J61196.AP, Thermo Fisher Scientific), 1×** for probe preparation, chip washing, baseline measurements, and target incubation.
**Tris(2-carboxyethyl)phosphine (TCEP; Pierce™ TCEP-HCl, No-Weigh™ Format, Catalog number A35349)** for reduction of disulfide-protected thiolated probes before immobilization.
**6-Mercapto-1-hexanol (MCH; 451088, Millipore Sigma)** for backfilling and passivation of the gold surface after probe immobilization.

● N-hydroxysulfosuccinimide (Sulfo-NHS; A3G26G, Thermo Fisher Scientific)

**1-ethyl-3-(3-dimethylaminopropyl) carbodiimide (EDC; A35391, Thermo Fisher Scientific)**
**P1 probe (P1; Integrated DNA Technologies; HPLC-purified):** thiolated 24-mer DNA probe labeled with a ferrocene redox reporter (P1: /5ThioMC6-D/TA CCA GCT ATT GTA TCT AAT AAG A/3FerrK/).
**P2 construct (P2; Integrated DNA Technologies; HPLC-purified):** For antibody- and nanobody-based sensors, P2 contains a distal carboxyl group at 5’ end for covalent coupling of the antibody or nanobody. P2 sequence complementary to P1: 5′-TCT TAT TAG ATA CAA TAG CTG GTA-3′.
**Antibody- or nanobody–P2 construct (Ab- or nb-P2):** antibody (IL-1 beta Monoclonal Antibody (2805), Thermo Fisher Scientific) or nanobody (Recombinant Anti-human TNF VHH Single Domain Antibody, Creative Biolab) that can conjugate to P2 sequence complementary to P1.
**Ethanolamine (411000; Sigma-Aldrich)**
**MES (MES Buffered Saline Pack; 28390, Thermo Fisher Scientific)**
**TM buffer (TM Buffer, 1X Tris-Magnesium sulphate, pH 7.5; RLMB-059, VWR)**
**Target proteins** for calibration and sensing experiments, prepared at the required concentrations in 1X PBS or other appropriate assay buffer. TNF-α (Human TNF-alpha Recombinant Protein, PeproTech®(300-01A), Thermo Fisher Scientific); IL-1β (Human IL-1 beta Recombinant Protein (PHC0816), Thermo Fisher Scientific)
**Nitrogen gas** for drying chips after cleaning and rinsing steps.

### Reagent setup

#### Gold electrodeposition solution

Prepare 50 mM HAuCl_4_ in 0.5 M HCl immediately before use or store under conditions recommended by the manufacturer. Protect from light where appropriate.

#### P1 reduction solution

Prepare two separate solutions containing 100 µM P1 and 100 mM TCEP in 1× PBS. Mix 2 µl of the 100 µM P1 with 2 µl 100 mM TCEP solution. Incubate in the dark for 1 h before hybridization or immobilization.

#### P1/P2 hybridization mixture for aptamer-based sensors

Mix 4 µl of reduced P1 with 96 µl of PBS. Then add 100 µl of 2 µM Apt-P2, reaching 200 µl in total volume. Hybridize using the defined thermal program before application to the chip.

#### MCH backfilling solutions

Prepare 1 mM and 10 mM MCH solutions in 1× PBS or in the buffer used routinely for monolayer formation. For each concentration, prepare 2 ml of solution immediately before use or from a freshly opened MCH stock bottle.

▲ **Critical step** MCH should be handled as a fresh reagent because thiol-containing compounds can oxidize during storage and repeated exposure to air. Minimize bottle opening time, close the bottle immediately. Use freshly prepared MCH working solutions for each experiment. Keep the MCH stock bottle at 4 °C and replace the bottle every 3 months, or sooner if inconsistent monolayer formation, increased background signal or poor sensor reproducibility is observed.

***▴***Caution

HAuCl_4_, HCl, and H_2_SO_4_ are corrosive and should be handled using appropriate PPE and approved chemical safety procedures.
Thiolated DNA probes and conjugated probe constructs should be handled in low-binding tubes and stored according to manufacturer or synthesis specifications to minimize degradation.
Ferrocene-labeled probes should be protected from prolonged light exposure and repeated freeze–thaw cycles where possible.
MCH has a strong odor and should be handled in a chemical hood.

Equipment

**Potentiostat/galvanostat** capable of CA, CV, and Square Wave Voltammetry (SWV), with sufficient temporal resolution and bandwidth for MP measurements. The instrument should be evaluated under the same current range and waveform used in the experiment.
**Three-electrode electrochemical setup**, including:
patterned Ti/Au chip as the working electrode,
Ag/AgCl reference electrode,
platinum wire counter electrode.
**Plasma cleaner** for chip surface cleaning before nanostructuring or functionalization.
**Micropipettes and low-retention pipette tips** for handling probe solutions, buffers, and target solutions.
**Humidified dark chamber** for overnight probe immobilization.
**Thermocycler or temperature-controlled heating block** for probe hybridization.
**Nitrogen gun or nitrogen line** for drying chips after rinsing.
**Chemical-resistant containers or reservoirs** for chip cleaning and rinsing.
**Electrochemical cell or chip holder** compatible with the patterned sensor chips and small-volume measurements.
**Analytical balance** for reagent preparation, if solutions are prepared from solid reagents.
**pH meter** for preparation and verification of buffer and acid solutions, where needed.
**Vortex mixer** for preparing homogeneous reagent solutions.
**Centrifuge / mini-centrifuge** for collecting small-volume probe solutions.
**Optical microscope** for visual inspection of chips before and after nanostructuring and functionalization.
**Refrigerator (4 °C) and freezer (−20 °C or −80 °C, as appropriate)** for storage of DNA probes, conjugates, and protein targets.

### Equipment setup Electrochemical workstation

Set up the electrochemical workstation before beginning any sensor measurements. Connect the working electrode (WE), reference electrode (RE), and counter electrode (CE) to the corresponding potentiostat leads according to the manufacturer’s configuration. For a conventional three-electrode setup, connect the MP sensor electrode to the WE lead, the Ag/AgCl or Ag reference electrode to the RE lead and the Pt wire, Pt mesh or other inert auxiliary electrode to the CE lead (See Box 5). If the potentiostat uses separate working-sense or reference-sense leads, connect these according to the manufacturer’s instructions and verify that all unused leads are properly parked or disconnected to avoid electrical noise.

Place the RE as close as practically possible to the WE without touching the sensing area, blocking mass transport or disturbing the solution meniscus. This minimizes uncompensated resistance and improves reproducibility of the applied potential. Position the CE so that it provides a stable current path through the solution but does not physically contact the WE, RE or sensor surface. The CE should have a surface area equal to or larger than that of the WE to avoid current limitation at the auxiliary electrode. Maintain the same WE–RE–CE geometry for all baseline, calibration and target measurements.

For chip-based measurements with patterned electrodes, connect the WE contact pad to the potentiostat using a probe station, spring-loaded pogo pin, microclip or conductive connector. Confirm that the connector contacts only the intended metal pad and does not bridge adjacent electrodes. If multiple WEs are present on the same chip, connect only the selected WE during individual measurements unless the potentiostat is configured for multiplexed operation. Keep the chip, electrode leads and connectors mechanically stable throughout the experiment, as small movements can introduce current spikes or baseline shifts.

Before measuring modified sensors, confirm that the open-circuit potential is stable, the background current is within the expected range, and the current response changes appropriately when the WE, RE, or CE is disconnected. A flat or saturated response often indicates an incorrect electrode connection, poor contact to the WE, a disconnected RE, a short circuit between electrodes or insufficient solution coverage.

For CA MP measurements, operate the potentiostat under conditions that provide sufficient bandwidth, sampling rate and current sensitivity to resolve rapid current transients reproducibly. Use the same current range, sampling interval, filtering, quiet time and acquisition window across all baseline, calibration and target measurements. For representative ferrocene-labelled MP measurements, apply a potential step from 0 to +500 mV versus Ag/AgCl and record the current transient over 50 ms (See CA parameters in Box 5). The acquisition settings should be optimized so that early-time current values, such as the current at 140 µs, can be extracted reliably without saturation, excessive filtering or undersampling.

▲ **Critical step** Incorrect electrode connection is one of the most common causes of failed electrochemical measurements. Reversing the WE and CE, using an unstable RE or allowing the RE to dry can generate distorted transients, unstable baselines or apparent sensor responses that are unrelated to target binding.

▲ **Critical step** The RE must remain immersed and stable throughout the measurement. Do not allow the reference junction to contact air, dry out or sit outside the solution droplet. For droplet-based measurements, ensure that the WE, RE, and CE are all immersed in the same continuous liquid volume.

▲ **Critical step** Avoid placing electrode leads near moving equipment, power supplies or unshielded cables. Use a Faraday cage, when possible, especially for low-current measurements or fast CA acquisition.

### Hybridization setup

Program the thermocycler or heating block for probe hybridization as follows: 95 °C for 5 min, followed by stepwise cooling to 55 °C for 2 min, 45 °C for 2 min, 35 °C for 2 min and 25 °C for 2 min. Use low-binding tubes and ensure that all oligonucleotide solutions are fully mixed and briefly centrifuged before heating. After hybridization, keep the probe solution protected from light if it contains ferrocene.

### Humidified incubation chamber

Prepare a light-protected humidified chamber before probe immobilization. Place clean, lint-free tissue or absorbent pads wetted with DI water or PBS inside a sealed container, ensuring that the liquid reservoir does not directly contact the sensor chip. Place the sensor chip on a clean, level support and apply the probe solution to the active electrode area. Close the chamber immediately to minimize evaporation during overnight immobilization.

▲ **Critical step** Evaporation during immobilization can concentrate salts and probes, alter surface assembly and produce nonuniform monolayers. The chamber should remain humidified and light-protected for the full immobilization period.

#### (B) Electrode preparation

**43. Fabrication of Nanostructured Microelectrodes (NMEs)**

Surface Preparation: Rinse sequentially with acetone, IPA, and DI water, then dry with nitrogen to remove the protective S1805 layer. Etch chips with plasma for 1 minute.

Gold Solution Preparation: Prepare 50 mM HAuCl_4_ in 0.5 M HCl.

Electrodeposition: Step 1 (Microstructure) – Using a three-electrode setup with an Ag/AgCl reference electrode and a Pt wire counter electrode, perform constant-potential electrodeposition in the gold electrodeposition solution. In the first deposition step, apply 0 mV versus Ag/AgCl for 100 s. This step generates the microscale three-dimensional gold structure, typically with a diameter of approximately 100 µm and wavy features of approximately 250 nm.

Step 2 (Nanostructure) – In the second deposition step, using the same solution and electrode configuration, apply −450 mV versus Ag/AgCl for 10 s. This step generates fine nanofeatures, typically <20 nm, that are important for high-sensitivity molecular pendulum sensing.

For Autolab instruments, select the chronoamperometry procedure from the default procedures and apply 0 V for 100 s, followed by −450 mV for 10 s. For BASi instruments, select DC potential amperometry (DCPA) from the New Experiment menu and apply 0 V for 100 s, followed by −450 mV for 10 s. These settings implement the same two-step constant-potential electrodeposition process, although the method name differs between instruments.

Record the current trace during both deposition steps for every electrode. The current profile provides a practical quality-control readout of electrode cleanliness, fabrication quality, exposed electroactive area and reproducibility of gold growth. For reproducible electrodes, deposition traces should have similar shapes and should reach comparable final current values at the end of the 100 s microstructure deposition step and the 10 s nanostructure deposition step. Large deviations in the current profile may indicate contamination, poor electrical contact, incorrect electrode area, incomplete passivation, or fabrication defects.

Quality Control: Wash with DI water and dry. Check the NME morphology and size under the microscope. Perform CV in 100 mM H_2_SO_4_ (0 to 1500 mV at 100 mV/s) for a minimum of 40 cycles to electrochemically clean gold and confirm stable successive voltammograms.

#### **(C)** Probe preparation

**44. Probe Preparation and Chip Functionalization for Aptamer-Based Sensors**

Reduction: Mix 100 µM P1 (2 µl) with 100 mM TCEP (2 µl) in PBS and incubate in the dark for 1 hour to reduce disulfides.

Hybridization: Mix 2 µM P1 (100 µl) and 2 µM P2 (100 µl), then incubate the mixture at 95 °C for 5 min, ramping down to 55 °C for 2 min, 45 °C for 2 min, 35 °C for 2 min, and 25 °C for 2 min.

Immobilization: Drop 20 µL of this solution onto the chip, ensuring NMEs are fully covered. Incubate overnight in a dark, humid chamber.

Backfilling: Drop 1 mM MCH overnight, followed by 10 mM MCH for 30 min. Washing: Wash three times for 5 minutes with 1X PBS.

**45. Probe preparation and chip functionalization for antibody-based and nanobody-based sensors**

Reduce the thiolated P1 strands in 1 mM TCEP by incubating the solution in the dark at room temperature for 1 h. Mix the reduced P1 strands with the P2-base strand containing the reactive carboxyl group and dilute the mixture in TM/PBS buffer to a final concentration of 1 µM. Heat the solution to 95 °C for 5 min and then cool it to 4 °C to form the P1–P2 duplex framework. Apply the pre-hybridized DNA framework to the gold electrodes and incubate overnight to allow immobilization. After incubation, rinse the electrodes thoroughly to remove unbound material with PBS. (Note: Incubation of antibody-containing sensors with TCEP can induce degradation and should be avoided. While protocols used in the past for MP sensors have included TCEP that was carried through to steps where contact with antibody receptors was possible, the reagent was stored in PBS, which strongly attenuates activity and suppresses antibody degradation. Other conjugation strategies can also be used to couple antibody or nanobody receptors to the DNA framework, depending on the receptor format, available functional groups and desired orientation.)

Activate the carboxyl groups on the immobilized P2 strands by incubating the electrodes in MES buffer (0.1 M) containing a 2:1 mixture of sulfo-NHS (100 mM) and EDC (50 mM) for 30 min. Following activation, incubate the electrodes with antibody/nanobody solution prepared in PBS at a final concentration of 200 ng ml^-1^ for 6 h to enable covalent conjugation. Rinse the electrodes with PBS to remove unreacted reagents and excess antibody/nanobody, then bundle the chips together before surface blocking.

▴**Critical step** Carbodiimide coupling is a well-established conjugation method that is widely used in biosensing and surface biofunctionalization. MES buffer is commonly used for EDC/sulfo-NHS activation because it supports efficient carboxyl activation while minimizing competing amine-containing species. However, if MES is not compatible with a specific receptor, surface chemistry, or downstream assay, PBS can be used as an alternative activation or conjugation buffer, provided that the incubation time is optimized and coupling efficiency is verified experimentally. Longer antibody or nanobody incubation times may improve surface coupling, but they can also increase nonspecific adsorption, receptor crowding, or loss of binding activity. Therefore, receptor concentration and incubation time should be optimized for each antibody or nanobody rather than treated as universal parameters.

Incubate the bundled electrodes overnight in 1 mM 6-mercapto-1-hexanol (MCH) prepared in PBS. Follow this with an additional incubation in 10 mM MCH for 30 min to ensure more complete passivation of exposed gold regions and to minimize nonspecific adsorption. Quench any remaining activated carboxyl groups by incubating the electrodes in 1 mM ethanolamine for 10 min. Wash the functionalized electrodes three times by immersion in 1× PBS for 5 min each before electrochemical characterization or target sensing.

#### **(D)** Analysis and sensing

**46. Probe Verification**

Perform SWV (0 to 500 mV, 60 Hz, and 100 nA current range). A distinct ferrocene peak at ∼380 mV vs Ag/AgCl indicates successful modification (See Box 5 for the SWV parameters and representative signal).

▲**Critical step** Ensure that the reference electrode (RE) and counter electrode (CE) are positioned reproducibly for all measurements. The RE should be placed as close as practically possible to the working electrode (WE), without touching the sensor surface or obstructing mass transport, to minimize uncompensated resistance and potential error. The CE should be positioned to provide a stable current path without physically perturbing the solution above the WE. Variations in RE/CE placement can change the effective potential experienced by the surface-bound probes and introduce apparent signal differences that are unrelated to target binding.

▲**Critical step** SWV should be used as a verification step, not as a repeated monitoring measurement. Repeated potential cycling can perturb the electrode interface, alter the organization of the thiol-based monolayer, promote partial reporter oxidation or loss, and reduce subsequent signal stability. Once a clear ferrocene peak has been confirmed, avoid repeating SWV on the same sensor unless the CA response is unreasonable, the baseline drift exceeds the acceptable threshold, or there is a specific need to troubleshoot probe immobilization or reporter accessibility. For routine sensing, use CA measurements after the initial SWV verification.

**47. Baseline Measurement**

Take an initial CA reading in 1× PBS using a potential step from 0 to +500 mV for 50 ms with 100 µA current range. After 30 min of incubation in 1× PBS, take a second CA reading under the same conditions to assess baseline stability. A signal change of more than 10% relative to the initial CA response should be considered drift.

A control sensor without a receptor, or a receptor-modified sensor maintained in buffer for the full experimental period, should be included to evaluate signal drift. If drift greater than 10% is observed, troubleshooting is required to assess monolayer stability, electrode contact, evaporation, reference-electrode stability, nonspecific interfacial changes and instrumental variability. In this case, subsequent signal changes should not be interpreted as specific binding unless the drift source is identified and controlled.

▲**Critical step** Maintain a constant solution volume and electrode geometry throughout baseline and target measurements. Even small changes in buffer droplet volume or electrode immersion depth can alter solution resistance, capacitive background and apparent faradaic current. This is especially important when measurements are performed in small-volume droplets or confined reservoirs.

▲**Critical step** Minimize evaporation during all calibration and sensing measurements. Evaporation increases the effective salt and protein concentrations, changes ionic strength, shifts interfacial properties and can cause artificial signal drift. For all measurements and incubations, use covered reservoirs or sealed chambers when possible, keep the sensor setup in a humidified environment and verify that the liquid volume remains constant over the full experiment.

▲**Critical step** The orientation of the sensing setup should be selected based on the measurement format. A vertical configuration, in which the sensor is immersed in a larger solution volume, is generally preferred when evaporation is a concern because it provides more stable hydration and reduces changes in droplet geometry. A horizontal configuration can be used for droplet-based measurements, but it should be performed in a humidified chamber and with careful control of droplet volume and meniscus position. Do not compare calibration data obtained from vertical and horizontal formats unless the geometry, volume and mass-transport conditions have been validated to be equivalent.

- **◆** Troubleshooting

**Problem:** The baseline signal increases during repeated chronoamperometric measurements.

**Possible reason:** Surface blocking with MCH is insufficient, or the monolayer contains defects that expose regions of bare or poorly passivated gold. These defects can increase nonspecific adsorption, alter interfacial capacitance and produce an apparent forward shift in the baseline.

**Solution:** Incubate the sensor in 10 mM MCH for 30 min to improve passivation of exposed gold regions. Rinse thoroughly with 1× PBS and repeat the baseline measurement. If the baseline remains unstable, prepare a new sensor and evaluate whether the immobilization or blocking steps require optimization.

**Problem:** The baseline signal decreases during repeated chronoamperometric measurements.

**Possible reason:** Washing steps may have been insufficient to remove excess or weakly adsorbed material from the electrode surface, or the coated monolayer may not have had sufficient time to stabilize before measurement. Gradual desorption, rearrangement or removal of loosely associated material can produce a backward shift in the baseline.

**Solution:** Replace the buffer with fresh 1× PBS and allow the sensor to equilibrate for at least 30 min. After equilibration, repeat the CA measurement several times under identical conditions to assess whether the baseline has stabilized. Do not proceed to target detection until the baseline drift is

≤10% relative to the initial CA response.

**48. Target Detection**

Incubate the sensor with a series of target protein concentrations selected according to the apparent binding affinity of the receptor–target pair and the intended analytical range. For initial calibration, prepare a serial dilution of the target protein in the assay buffer, typically spanning approximately tenfold below to tenfold above the apparent dissociation constant, K_D_, of the recognition element. This range can be adjusted after the first calibration experiment if the response saturates too early, remains near baseline across the tested range or does not cover the expected physiological concentration window. Representative affinity-dependent concentration ranges for TNF-α and IL-1β, together with the corresponding bio-layer interferometry-derived K_D_ values, are provided in Box 4.

Expose the sensor sequentially to increasing concentrations of target protein. Use the same incubation volume, electrode configuration and temperature for all concentrations. Incubate the sensor for 20–60 min at each concentration, or until the electrochemical response approaches a stable value. The required incubation time should be determined empirically for each receptor– target pair, as it depends on receptor affinity, surface density, target size, diffusion, transport within the measurement geometry and association kinetics. For the first calibration experiment, use a fixed incubation time across all concentrations to avoid introducing time-dependent bias. Once the binding kinetics are known, the incubation time can be shortened or extended, provided that the same timing is used consistently within a given calibration series.

After each incubation step, record a CA trace using the same potential step and acquisition settings used for the baseline measurement. For example, apply a potential step from 0 to +500 mV versus Ag/AgCl for 50 ms and record the ferrocene-mediated current response. Avoid changing the current range, sampling rate, filtering or acquisition window between concentrations, as these parameters can alter the apparent current decay profile. If solution exchange is performed between concentrations, remove and replace the solution carefully without exposing the sensor surface to air, damaging the gold nanostructure, or disturbing the position of the working, reference or counter electrodes.

For cumulative titration experiments, add target solutions sequentially from low to high concentration and record the response after each incubation period. This format minimizes sensor-to-sensor variability and is useful for establishing the concentration-response relationship on the same modified surface. However, because cumulative titration does not fully remove previously bound targets, it should be interpreted as an apparent binding curve unless an effective regeneration step is included between concentrations. For independent-concentration calibration, use separate sensors for each target concentration or regenerate the same sensor between measurements only if regeneration has been validated not to damage the monolayer, receptor or electrochemical reporter. Independent-concentration measurements are preferred when assessing reversibility, hysteresis or possible carryover.

Include at least one buffer-only control sensor that is exposed to the assay buffer for the full duration of the experiment. This control is used to quantify baseline drift arising from monolayer relaxation, reporter instability, nonspecific interfacial changes, evaporation or instrumental drift. In addition, include a non-binding control sensor, such as a sensor lacking the recognition receptor or carrying a scrambled/non-specific receptor, when evaluating new protein targets or complex matrices. The target-associated response should exceed the drift observed in the buffer-only control and the nonspecific response observed in the non-binding control. If the control sensors show concentration-dependent changes comparable to the receptor-modified sensor, the response should not be assigned to specific target recognition.

Extract the current at a fixed time point, such as 140 µs, to generate calibration curves. The signal-extraction time point should be selected based on the potentiostat bandwidth, sampling resolution, RC charging behavior, electrode geometry and electron-transfer kinetics of the reporter, and must be kept constant across all concentrations, replicates and controls within a given calibration experiment. When appropriate, CA traces may be normalized, for example, by setting the maximum current within each trace to 1, to facilitate comparison of transient shape or relative signal change. However, normalization is not required for all analyses, and raw or baseline-referenced currents may be more appropriate when absolute current magnitude, electrode-to-electrode variability or signal amplitude is being evaluated. Plot the selected signal output, such as raw current, normalized current, baseline-referenced current or percent signal change relative to the pre-target baseline, as a function of target concentration. Fit the resulting curve using an appropriate binding or empirical calibration model only after confirming that the baseline and control responses remain stable over the full measurement period.

Perform calibration measurements using replicate sensors and, when possible, independently prepared protein dilution series. A minimum of three independently prepared sensors is recommended for initial calibration of a new receptor–target pair. Technical replicates from the same protein dilution series are useful for assessing measurement precision, but they should not be treated as fully independent biological or sensor-fabrication replicates. Report the number of independent sensors, the number of repeated measurements per sensor, the concentration sequence and whether the calibration was performed by cumulative titration or independent-concentration exposure.

▴**Critical step** Protein solutions at low concentration are susceptible to loss through degradation, aggregation or adsorption to the inner surfaces of tubes, pipette tips, wells and reservoirs. Prepare low-concentration standards immediately before use, use low-binding plasticware where appropriate, avoid unnecessary transfers and maintain consistent incubation times. Apparent decreases in faradaic signal at low target concentrations often reflect analyte loss or nonspecific adsorption rather than true sensor performance.

▲**Critical step** Avoid introducing bubbles near the WE, RE, or CE during solution exchange. Bubbles can partially block the active sensing area, disrupt the current path or alter local mass transport, producing abrupt changes in CA traces. If bubbles are observed, remove them before acquiring electrochemical data.

▲**Critical step** Keep the temperature constant during calibration. Protein binding kinetics, diffusion, solution resistance and ferrocene electron-transfer behavior are temperature-sensitive. Temperature fluctuations during sequential concentration measurements can therefore appear as false concentration-dependent responses.

▲**Critical step** Do not compare calibration curves acquired using different incubation formats, measurement geometries or solution volumes unless these conditions have been validated to produce equivalent mass transport and baseline stability. Differences in droplet height, electrode spacing, evaporation rate or convective mixing can change the apparent response even when the receptor–target interaction is unchanged.

▲**Critical step** For cumulative titrations, always proceed from the lowest to the highest target concentration. Reversing the order can produce carryover from high-concentration exposure and obscure the response at lower concentrations. If the sensor is intended to be reused after high-concentration exposure, regeneration efficiency and post-regeneration baseline recovery must be validated separately.

##### BOX 5 Equipment setup, SWV, and CA parameters along with ferrocene peak

**Fig. 1.**
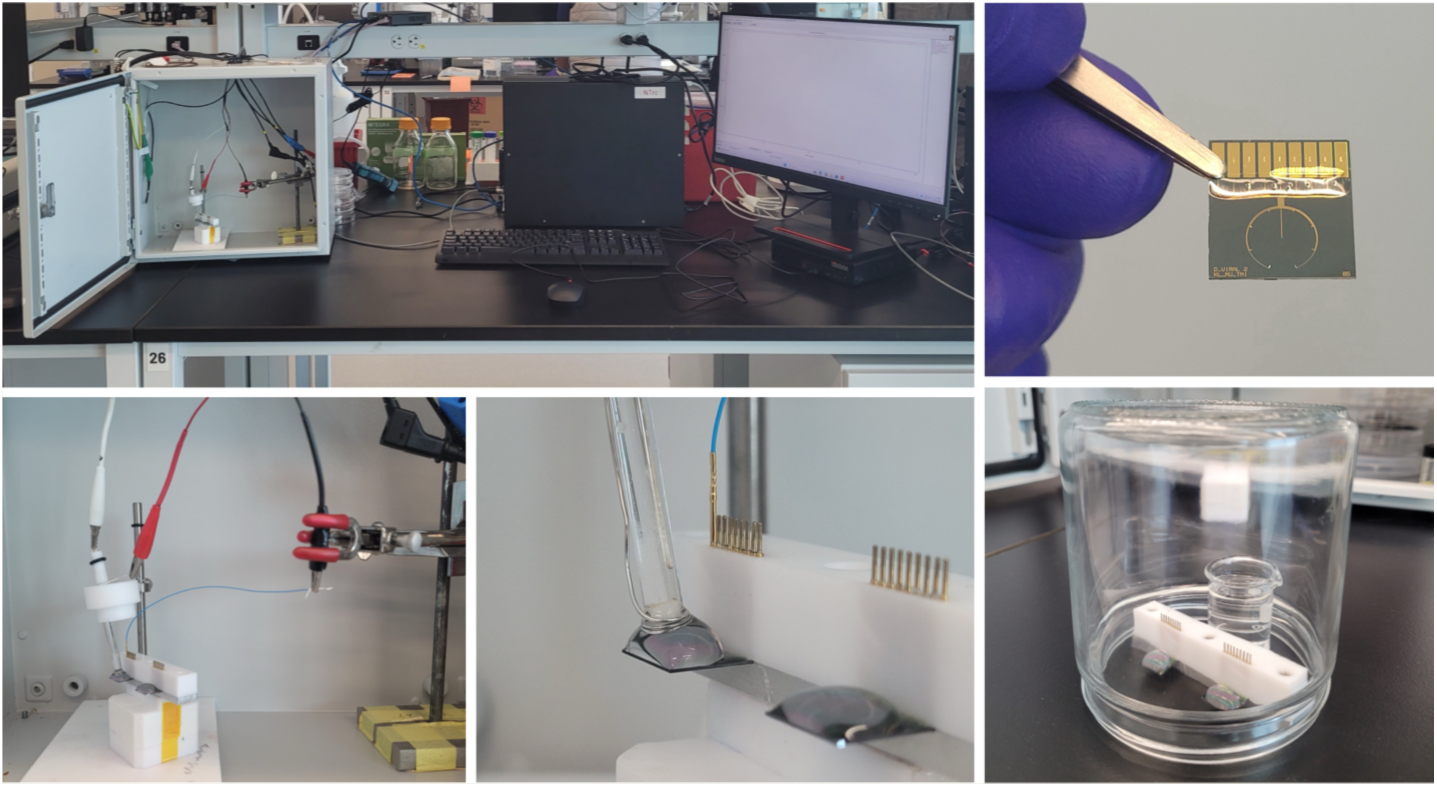
Experiment setup for electrochemical measurement along with incubation chamber

**Fig. 2.**
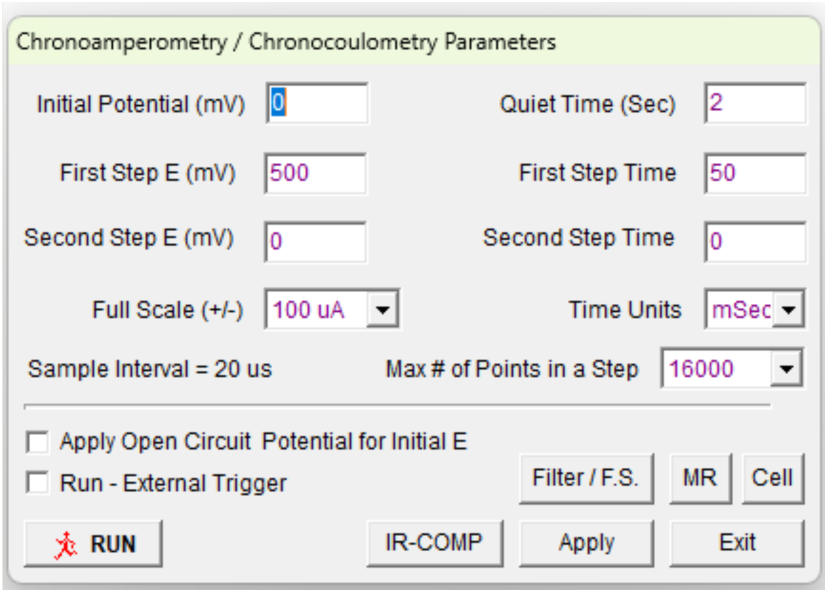
CA parameters for measurements in BASi Epsilon potentiostat

**Fig. 3.**
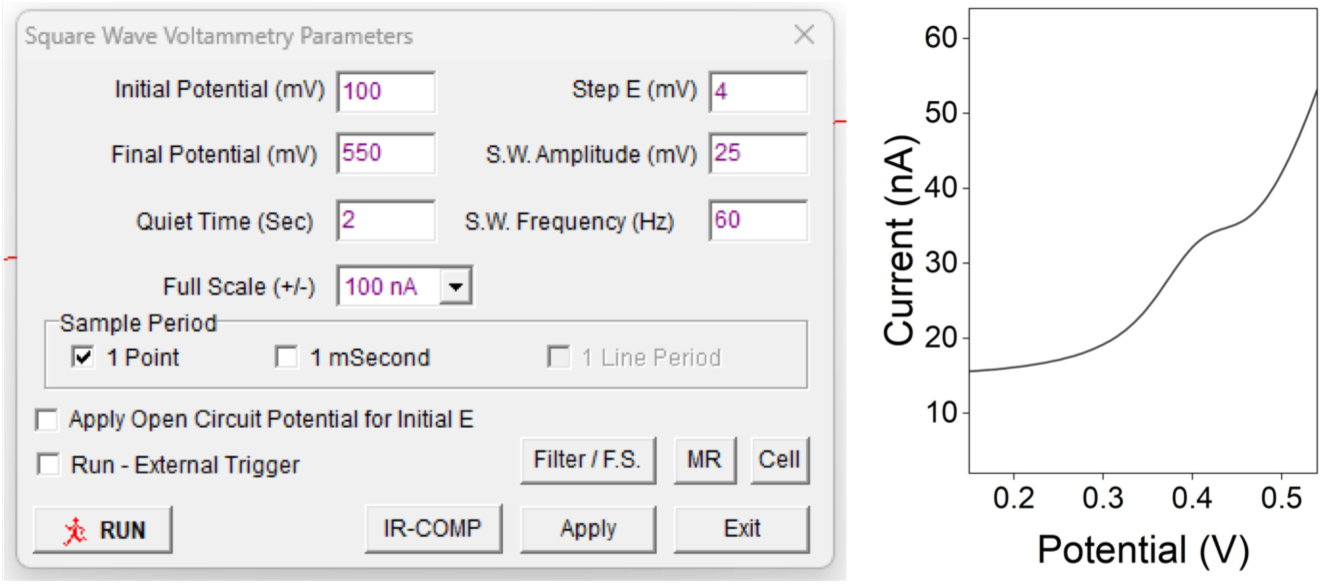
SWV parameters in BASi Epsilon potentiostat for probe existence evaluation along with Ferrocene peak vs. Ag/AgCl

### Active-Reset Operation G Parameters

Active-reset is used to regenerate the MP sensor after target binding by applying an oscillating electrical waveform that accelerates dissociation and restores the baseline sensing state. Because reset efficiency depends strongly on waveform conditions, the voltage amplitude and oscillation frequency must be optimized empirically for each receptor–target system and measurement configuration. The goal is to identify conditions that maximize signal recovery after binding while minimizing probe degradation, baseline drift, monolayer disruption, and loss of sensing performance over repeated cycles.

#### **(A)** Experimental design

The optimization procedure should be performed on fully functionalized sensors after target binding has produced a stable bound-state signal. Reset conditions are then screened systematically by varying voltage amplitude and frequency while keeping all other experimental variables constant, including electrolyte composition, temperature, electrode geometry, target concentration, incubation time, and readout waveform. After each reset condition is applied, the sensor is interrogated again using the standard CA sensing method, and the extent of signal recovery is quantified relative to the initial unbound baseline and the bound-state signal. The optimal reset condition is the one that produces rapid and reproducible recovery of the unbound-state signal without introducing progressive signal decay, increased noise, or changes in probe behavior over repeated cycles.

Reagents

Fully functionalized MP sensor chips
Target protein solution at a concentration sufficient to generate a stable bound-state signal
1× PBS or the standard sensing buffer used for the assay
DI water for rinsing, if needed
Nitrogen gas for drying, if needed

Equipment

Potentiostat capable of applying user-defined oscillating potential waveforms with sufficient temporal resolution (e.g., Metrohm Autolab PGSTAT302N)
Three-electrode electrochemical setup, including:
MP-modified sensor chip as the working electrode
Ag/AgCl reference electrode
platinum wire counter electrode
Electrochemical cell or chip holder compatible with small-volume measurements
Micropipettes and low-retention pipette tips
Timer
Faraday cage or equivalent shielding setup, if used for sensing measurements

#### (B) Procedure

**49. Establish the baseline and bound-state signals**

i. Place the functionalized sensor in the electrochemical cell containing 1× PBS or the sensing buffer.
ii. Record the baseline unbound-state CA response using the standard sensing waveform and acquisition settings.
iii. Incubate the sensor with the target protein under the predefined assay conditions until a stable bound-state signal is obtained.
iv. Record the bound-state CA response.

▲**Critical step** The baseline and bound-state signals must be collected using exactly the same readout conditions that will later be used to assess reset performance. Changing measurement parameters during optimization makes the comparison useless.

**50. Define the reset waveform screening matrix**

i. Select a range of reset voltage amplitudes to test.
ii. Select a range of oscillation frequencies to test.
iii. Keep all other waveform parameters fixed during the initial screen, including total reset duration, duty cycle, and number of cycles, unless those variables are themselves being optimized in a secondary screen.

A practical initial screening strategy is to vary one parameter at a time or to use a small voltage– frequency matrix. Voltage amplitude is typically screened first at a fixed intermediate frequency, followed by frequency optimization at the best-performing voltage.

▲**Critical step** Do not begin with excessively aggressive reset conditions. High voltages or frequencies can damage the monolayer, perturb the probe architecture, or introduce irreversible signal loss.

**51. Apply the reset waveform**

i. After recording the bound-state signal, replace the solution with a fresh sensing buffer if required by the assay design.
ii. Apply the selected oscillating reset waveform to the sensor for the predefined reset duration.
iii. Allow the sensor to re-equilibrate in the sensing buffer for approximately 5 min after the reset step, if needed, before recording the next baseline or target-associated measurement.
iv. Record the post-reset CA response using the standard sensing waveform.

▴**Critical step** The reset waveform must be delivered under the same electrode configuration and shielding conditions used for sensing. Hardware-dependent differences in waveform fidelity can otherwise confound interpretation of reset efficiency.

*▲***Caution** If large baseline shifts, abnormal noise, distorted transients, or loss of ferrocene signal are observed immediately after reset, the selected condition may be too harsh and should not be continued.

**52. Quantify reset efficiency**

i. For each reset condition, compare the post-reset signal with:
ii. the initial baseline unbound-state signal, and
iii. the pre-reset bound-state signal.
iv. Calculate reset efficiency using a normalized recovery metric, such as the fraction of the bound-induced signal change that is recovered after reset.
v. Repeat the same reset condition on replicate sensors or across repeated cycles on the same sensor to assess reproducibility.

▲**Critical step** The signal metric used for this calculation, such as current at a defined time point, integrated transient area, or fitted transient parameter, should remain constant throughout the optimization study.

**53. Screen voltage amplitude**

i. At a fixed oscillation frequency, apply reset waveforms across the selected voltage range.
ii. After each reset, measure the recovery signal and quantify reset efficiency.
iii. Identify the voltage range that provides substantial recovery without progressive signal degradation.

In many cases, increasing voltage improves reset efficiency up to a point, after which surface damage, probe destabilization, or baseline distortion begin to dominate.

▲**Critical step** The optimal voltage is not necessarily the one giving the most immediate recovery in a single cycle. Priority should be given to conditions that remain stable over repeated operation.

**54. Screen frequency**

i. Using the selected voltage amplitude, repeat the reset process across the chosen frequency range.
ii. Quantify signal recovery, reproducibility, and baseline stability after each condition.
iii. Identify the frequency that provides the best balance between rapid regeneration and preservation of sensor integrity.

Frequency affects the field-driven dynamics of the MP structure and the efficiency with which the system is driven away from the bound state. Frequencies that are too low may be insufficient for effective reset, whereas frequencies that are too high may reduce waveform fidelity, introduce hardware limitations, or impose unnecessary stress on the sensing layer.

**55. Validate the final reset condition over repeated cycles**

i. Apply the selected voltage and frequency combination in repeated sensing-reset cycles on the same sensor.
ii. After each cycle, record:
iii. the bound-state response,
iv. the post-reset response,
v. baseline drift,
vi. signal-to-noise ratio,
vii. and any change in transient shape.
viii. Determine whether the selected reset condition supports stable and repeatable long-term operation.

▲**Critical step** A condition that performs well in one reset event but causes cumulative degradation over multiple cycles is not suitable for continuous-monitoring applications.

#### (C) Data analysis

Reset optimization should be evaluated using more than one performance metric. Recommended outputs include:

reset efficiency after each waveform condition,
time required for recovery,
signal drift across repeated cycles,
change in peak or normalized transient shape,
cycle-to-cycle coefficient of variation,
and retention of dynamic sensing range after repeated resetting.

Where possible, representative raw CA traces should be shown together with summary plots of reset efficiency as a function of voltage and frequency.

### Anticipated results

Successful optimization should identify a reset waveform that restores the post-binding signal toward the initial baseline while preserving the integrity of the sensing layer over multiple cycles. Under suboptimal conditions, reset may be incomplete, producing only partial recovery of the baseline. Under overly aggressive conditions, the sensor may show increased noise, diminished ferrocene response, altered transient shape, or irreversible signal loss.

- **◆** Troubleshooting

**Problem:** Poor recovery after reset

**Possible reason:** Voltage amplitude and/or frequency are not in the optimized range; reset duration too short; target dissociation too slow under tested conditions

**Solution:** Increase voltage or frequency gradually, or extend reset duration while monitoring for degradation

**Problem:** Large baseline shift after reset

**Possible reason:** Excessively harsh waveform; monolayer perturbation; probe desorption

**Solution:** Reduce voltage amplitude, shorten reset duration, or reduce the number of oscillation cycles

**Problem:** Increased noise or distorted transients after reset

**Possible reason:** Instrument bandwidth limitation, shielding problem, or waveform instability

**Solution:** Verify potentiostat performance under the reset conditions, improve shielding and grounding, and inspect all electrode connections. If needed, confirm waveform fidelity using an electrical dummy load, such as a resistor, capacitor or RC circuit with known values, connected in place of the sensor. The dummy load allows the applied waveform and current response to be tested without contributions from monolayer instability, target binding or electrode-surface chemistry

**Problem:** Good initial reset but progressive signal loss over repeated cycles

**Possible reason:** Gradual damage to the sensing interface or incomplete equilibration between cycles

**Solution:** Select a less aggressive reset condition and include a standardized equilibration step before each post-reset measurement

## Data availability

The main data supporting this protocol are available in our previous publications^2,8^ and can be obtained from the corresponding author upon reasonable request. Source data are provided with this paper.

## Acknowledgments

We thank the Biohub Chicago operations team for administrative and logistical support, and the Process and Product Development team at Tyndall National Institute for silicon-based device fabrication. This work was funded by Biohub.

## Author contributions

H.Z. and S.O.K. conceived the idea. V. J., Z. C., R. Q., L. L., S. K., F. E., and W. B. developed the protocol and performed the fabrication and testing. V. J., Z. C., R. Q., L. L., S. K., F. E., W. B., A. N., J. D., L. F. A. C., R. N., A. J. H. S., K. R., M. D. C., and J. D drafted the manuscript with input from S.O.K. and H.Z. All authors read, edited and approved the final manuscript.

